# Flexible client-dependent cages in the assembly landscape of the periplasmic protease-chaperone DegP

**DOI:** 10.1101/2022.10.17.512556

**Authors:** Robert W. Harkness, Zev A. Ripstein, Justin M. Di Trani, Lewis E. Kay

## Abstract

The periplasmic protein DegP, that is implicated in virulence factor transport leading to pathogenicity, is a bi-functional protease and chaperone that maintains protein homeostasis in gram-negative bacteria. To perform these functions, DegP captures clients inside cage-like structures, which we have recently shown to form through the reorganization of high-order preformed apo-oligomers, consisting of trimeric building blocks, that are structurally distinct from client-bound cages. Our previous studies suggested that these apo oligomers may allow DegP to encapsulate clients of various sizes under protein folding stresses by forming cage ensembles that can include extremely large cage particles. To explore the relation between cage and substrate sizes, we engineered a series of DegP clients of increasing hydrodynamic radii and analyzed their influence on DegP cage formation. We used dynamic light scattering and cryogenic electron microscopy to characterize the hydrodynamic properties and structures of the DegP cages that are adopted in response to each client. We present a series of flexible cage structures including novel 30mer and 60mer particles. Key interactions between DegP trimers and the bound clients that stabilize the cage assemblies and prime the clients for catalysis are revealed. We also provide evidence that DegP can form cages which approach subcellular organelles in terms of size.

**Significance statement:** Gram-negative pathogens export virulence factors that interfere with the function of host cells. This process is mediated by DegP, a protein which controls protein homeostasis in the periplasm of these bacteria and thus is a target for the development of novel antibiotics. DegP operates by incorporating client proteins inside cage-like structures to either recycle them or protect them from aggregation. Using a combination of dynamic light scattering measurements and cryogenic electron microscopy, we have shown that DegP can adopt many types of cages, some as large as subcellular organelles, depending on the size of the engaged client. This property likely enables DegP to capture different sized clients in response to protein misfolding stresses.

## Introduction

The cellular protein homeostasis network ensures that proteins achieve their native structure and are localized and degraded in a tightly regulated manner (1, 2). Defects in this quality control mechanism lead to the misfolding and aggregation of protein molecules, and, ultimately, to cell death (3). To promote the proper folding and recycling of proteins, cells express an array of protein chaperones and proteases (4). Some proteins have both functionalities, thus providing client substrates with the chance to refold prior to being recycled if refolding is unproductive. For example, members of the widely conserved family of <u>H</u>igh temperature <u>r</u>equirement <u>A</u> (HtrA) proteins (5–7) operate as bi-functional proteases and chaperones that are critical to maintaining cellular fitness and are additionally involved in cell motility, division, and programmed cell death (8).

DegP is one of the bacterial orthologues of the HtrA protein family that includes members of the antibiotic resistant ESKAPE pathogens which have been declared a global threat to human health (9). It operates within the periplasmic space of gram-negative bacteria, becoming overexpressed in response to a variety of cellular stressors such as heat (10), oxidative (11), and osmotic shock (12), and DegP-null cell lines are growth defective under stressed conditions (13), underscoring DegP’s critical protective function. In conjunction with its role in general periplasmic protein quality control, DegP has been implicated as major a contributor to bacterial pathogenicity due to its participation in shuttling virulence factors to the outer membrane (14–17) where they can be embedded or efficiently secreted to the extracellular space to interfere with the function of neighboring host cells. Clients of DegP include outer membrane proteins (OMPs) (18) and autotransporters (15, 19, 20) which are collectively involved in nutrient uptake, toxin export, host immune system evasion, and adhesion, among other virulence functions (15, 21, 22). The virulence-promoting aspect of DegP function makes it an important target for the development of novel antibiotics, as drugs that modulate DegP’s interactions with clients could restrict the severity of bacterial infections and promote clearance by the human immune system (23).

Structurally, mature DegP monomers (lacking the N-terminal periplasmic localization sequence that is cleaved upon entry to the periplasm) contain a serine protease domain followed by two PDZ domains (Fig. 1A top) (6). Three monomers become tightly associated via protease:protease domain interactions to yield a trimeric building block as the basic DegP functional unit with the PDZ1 and PDZ2 domains of each monomer oriented towards the trimer exterior (Fig. 1A) (6). Long loops within the protease domains, in addition to flexible linkers connecting the protease-PDZ1 and PDZ1-PDZ2 domains, provide DegP with a remarkable structural plasticity that facilitates the formation of higher-order oligomeric states which have been implicated in DegP function (Fig. 1A bottom) (24–26). Classically, the DegP structure-function paradigm was interpreted in terms of structural transitions between an apo resting hexamer state and substrate engaged cage-like oligomers featuring 12 or 24 monomers that are catalytically active (27). The resting hexamer structure, solved by X-ray crystallography (6), was found to form through inter-trimer protease:protease interactions and PDZ1^*i*^:PDZ1^*j*^ domain interactions (in what follows we will distinguish domains from separate trimers by the subscripts *i* and *j, i*≠*j*). In this state, the PDZ1 and protease domain binding sites are occluded, thus mitigating the unwanted access and subsequent proteolysis of client proteins. An additional “open” hexamer structure was also solved, which was thought to allow for initial substrate engagement (6). Subsequent structural studies revealed that client binding to the PDZ1 and protease domains causes the reassembly of DegP trimers into cage-like 12mer and 24mer structures that are mediated by PDZ1^*i*^:PDZ2^*j*^ interactions (Fig. 1B) (18, 24, 25, 27). These forms of DegP have been shown to proteolyze clients (27) yet they can also chaperone OMPs (18). The interplay between these two contrasting DegP functions is not currently well understood.

**Figure 1.**
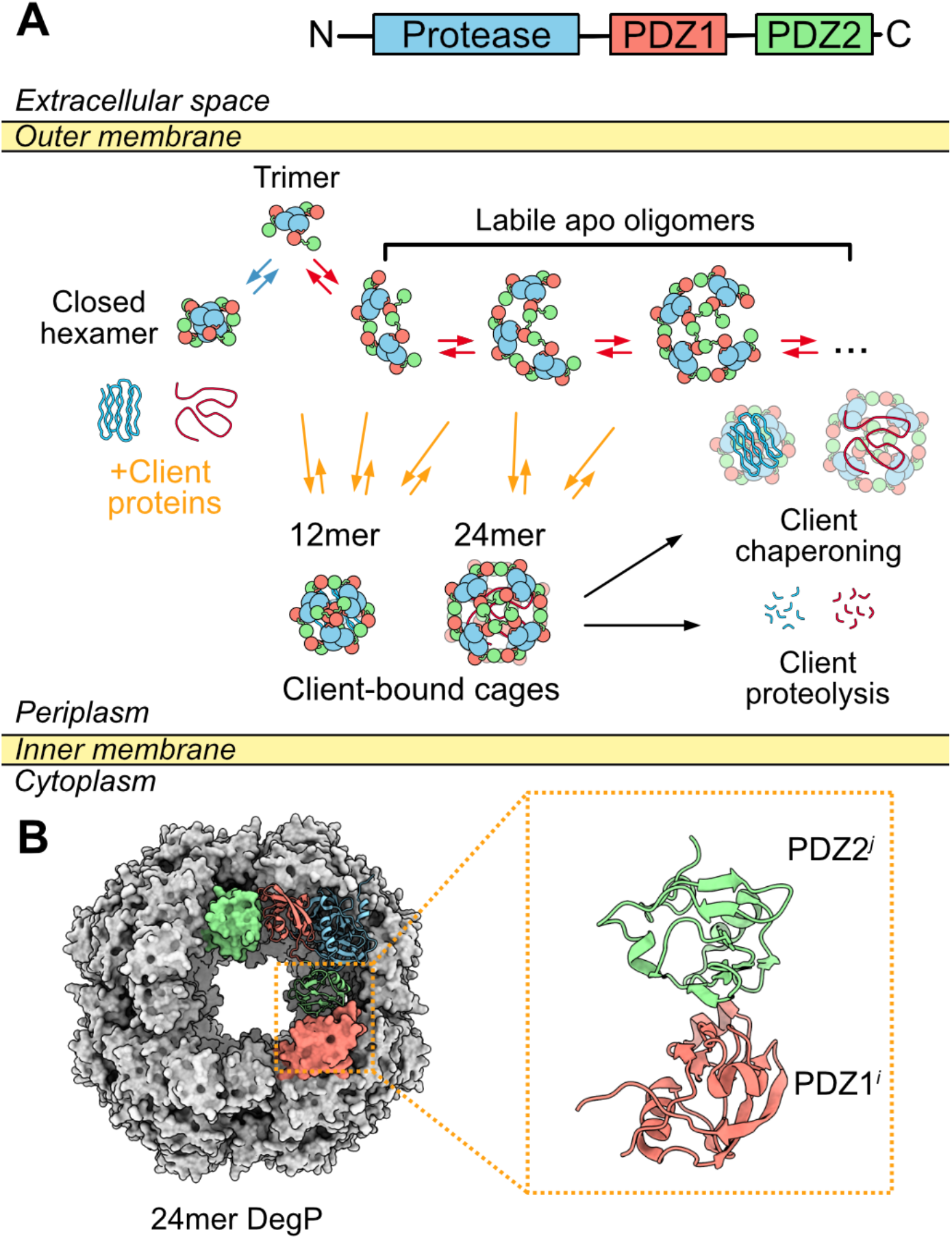
DegP forms client-dependent higher-order oligomers through inter-trimer PDZ domain interactions. (A) Mature DegP monomers contain a serine protease domain (blue) followed by two PDZ domains (PDZ1 in orange and PDZ2 in green, top). Three monomers associate via protease domain interactions to form trimeric building blocks that give rise to higher-order oligomeric architectures through inter-trimer PDZ1^*i*^:PDZ2^*j*^ domain interactions (bottom) (18, 25, 28). In the absence of substrates, DegP trimers assemble into a broadly populated ensemble of higher-order structures that are structurally distinct from bound cage conformations (28). Upon substrate engagement these apo oligomers reorganize into discrete cages, such as the 12mer or 24mer depicted here in cartoon form, which encapsulate the client (18). (B) Crystal structure of the octahedral 24mer cage (PDB 3MH6 (24), shown in grey surface representation) viewed down the 4-fold axis. Four of the eight trimers comprising the 24mer cage can be seen in the foreground of the image. A single protomer from the top right trimer, colored according to (A), is shown in ribbon representation, highlighting the relative protease/PDZ1/PDZ2 domain orientations in the context of the 24mer cage structure. This protomer is stabilized by an interaction between its PDZ2 domain (green ribbon) and an adjacent PDZ1 domain from a protomer in a second trimer located on the bottom right (shown as an orange surface), referred to as a PDZ1^*i*^:PDZ2^*j*^ interaction (enlarged to the right). A second such interaction is formed involving the PDZ1 domain from the initial protomer (red ribbon) and an adjacent PDZ2 domain (green surface) from a protomer in a third trimer (top left). This series of PDZ1^*i*^:PDZ2^*j*^ interfaces continues along the front face of the 24mer, returning to the initial interaction site after 4 successive contacts.

Recently, we have shown that in solution and in the absence of substrates, DegP populates a rapidly interconverting stress-dependent distribution of oligomeric states mediated by the weak self-assembly of trimers (Fig. 1A bottom) (28). The assembly landscape could be described in terms of two competing oligomerization pathways, one of which leads to the formation of the canonical hexameric state, and the other giving rise to a broadly populated ensemble of oligomers that are distinct from canonical, substrate-bound, cage conformers. Our data suggested that the ability to form this highly dynamic oligomeric ensemble comprised of partly-structured assemblies allows DegP to quickly respond to biological insults by engaging client proteins and reassembling into discrete cage conformers for substrate processing. Interestingly, our prior work implied that in the apo state, oligomeric ensembles of DegP can have effective molecular weights exceeding ∼3 MDa, corresponding to >60mer particles that are much larger than any known cage structures. We thus asked whether DegP actually forms cages of this size. Such large cage conformers may be of biological importance, for example, in the chaperoning of high molecular weight clients including autotransporter virulence factors, which are often ∼100-200 kDa in size (22, 29, 30). Furthermore, DegP likely encounters substrates of many sizes in the periplasm, and the ability to form a continuum of cage conformations, both apo- and client-bound, would enable a functional flexibility for responding to protein misfolding stresses. Here we have explored the types of cages that are adopted by DegP in response to substrates of different sizes. This was achieved through a recombinant client protein engineering approach in which we attached a known DegP binding motif to the C-terminus of a series of folded proteins of increasing hydrodynamic radii. We then analyzed the types of DegP cages formed in the presence of these chimeric clients using a combination of dynamic light scattering (DLS) and cryogenic electron microscopy (cryo-EM). We find a correlation between the size distribution of cage particles adopted by DegP and the size of the engaged substrate protein and herein present a series of client engaged structures including 30mer and 60mer cages. These reveal novel interactions between DegP trimers and bound clients that stabilize DegP cages and prime substrates for cleavage by the protease domains. We additionally provide evidence that, in the presence of large substrate proteins, DegP can form oligomeric structures that approach subcellular organelles in terms of size.

## Results

### Characterizing engineered DegP substrates and their influence on DegP cage distributions by DLS

DegP engages partially or fully unfolded client proteins via their C-termini (31), leaving their N-terminal portions unbound and, presumably, projected toward the interior of the cages. The unbound portions of clients are typically not observed in structures of DegP cages due to their flexibility. We hypothesized that by increasing the size of the N-terminal, free portion of the engaged substrate, DegP would be forced to adopt increasingly larger cages in order to accommodate clients within the cage interiors. We thus selected a series of folded proteins (TrpCage, His-SUMO, IL6, and MNeon) of increasing hydrodynamic radii (*r*_*h*_) to explore the influence of substrate size on the types of cages adopted by DegP (Fig. 2A). To ensure binding to DegP and provide each substrate with a standard interaction motif, each of these proteins was engineered to contain a known C-terminal DegP affinity tag. Here we selected the DNA binding domain of the human telomere repeat binding factor (hTRF1, 54 residues) (32) since we have previously shown it to interact with DegP with high affinity (*i*.*e*., complete binding saturation at ∼1:1 protomer:substrate molar ratio) (28). We refer to these engineered chimeric constructs, for example, as TrpCage-hTRF1 to denote their fusion to the C-terminal hTRF1 tag (see *SI Appendix*, Table S1 for their amino acid sequences).

**Figure 2.**
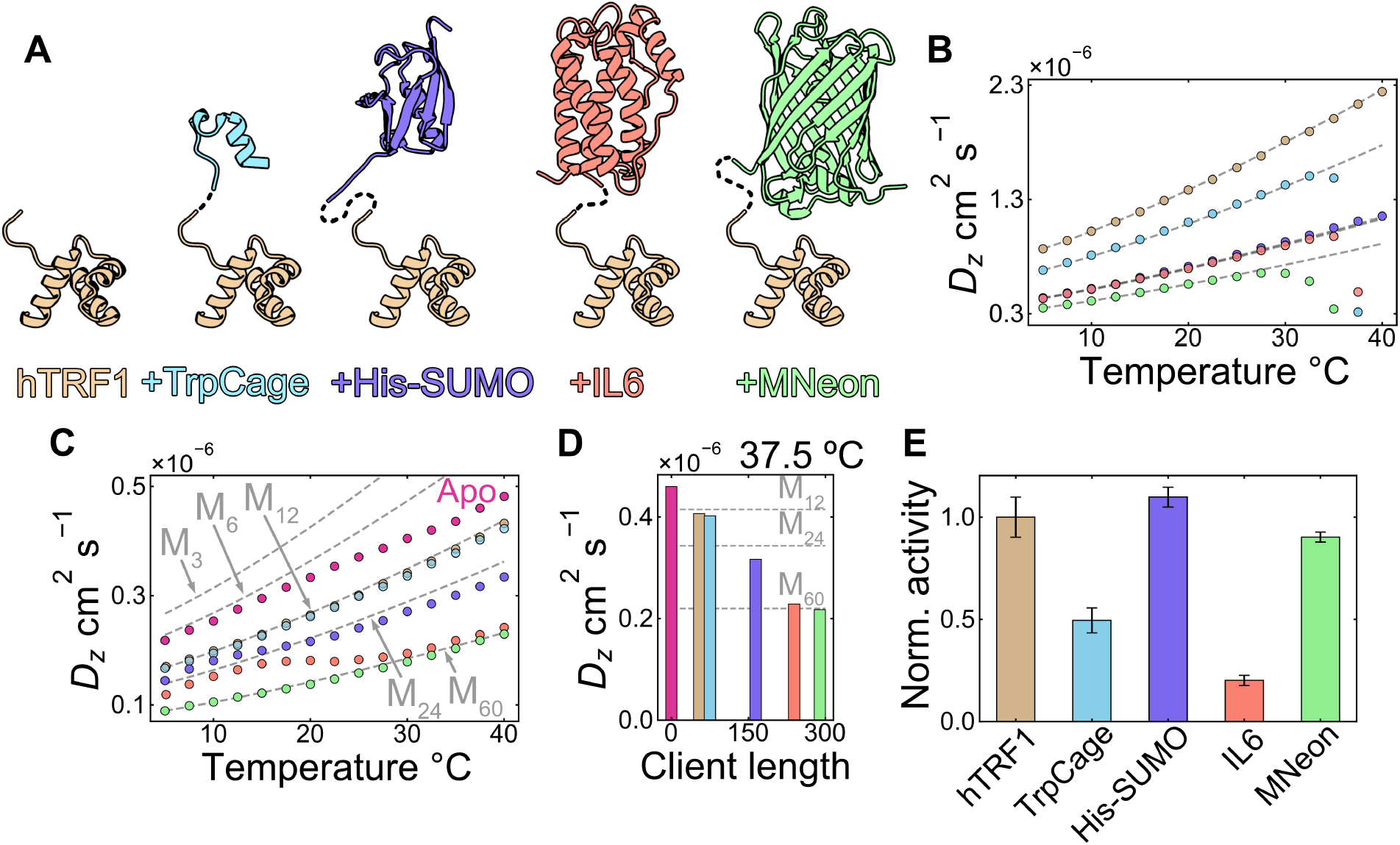
Characterization of engineered DegP clients and their influence on DegP oligomerization by DLS. (A) DegP substrates have been engineered with an hTRF1 binding tag (tan, PDB 1ITY (44)) C-terminal to folded proteins of increasing hydrodynamic radii (TrpCage, PDB 1L2Y (45), His-SUMO, PDB 1EUV (46), IL6, PDB 2IL6 (47), and MNeon, PDB 5LTR, in blue, purple, orange, and green respectively). (B) *D*_*z*_ values (colored points) for the substrates in (A) measured as a function of temperature by DLS. Note that certain substrates (TrpCage, IL6, MNeon) undergo irreversible unfolding/aggregation at high temperature (∼30-40 °C), leading to sharp decreases in measured *D*_*z*_ values. Hydrodynamic radii (*r*_*h*_) for each substrate (1.5, 1.9, 3.0, 3.0, and 3.7 nm respectively) have been calculated from their *D*_*z*_ values at 5 °C (in the absence of unfolding/aggregation) and used to generate temperature dependent *D*_*0*_ profiles to guide the eye (grey dashed lines), using the Stokes-Einstein relationship. (C) *D*_*z*_ values for S210A DegP (100 μM protomer concentration) in the presence of each of the substrates in (A) (200 μM). *D*_*z*_ values for apo DegP (no added substrate) are shown in red for comparison. *D*_*0*_ values (grey dashed lines) calculated for 3mer, 6mer, 12mer, 24mer, and 60mer DegP particles are indicated. In the case of the 3mer, 6mer, and 12mer, *r*_*h*_s of 4.9, 5.7, and 7.8 nm, respectively, were obtained from *D*_*z*_ values measured at low temperature (28), and then used to generate the *D*_*0*_(T) profiles shown via the Stokes-Einstein equation. Profiles for the 24mer and 60mer oligomers where established from values of *r*_*h*_s, 9.4 and 14.7 nm, respectively, as estimated at 20 °C using the structures determined here and the program HYDROPRO (49). (D) *D*_*z*_ values from (C) at 37.5 °C as a function of the client polypeptide chain length. *D*_*0*_ values for 12mer, 24mer, and 60mer particles are shown as horizontal grey dashed lines. (E) Normalized cleavage rates for the substrates in (A) in the presence of proteolytically active DegP. The cleavage activities for the engineered substrates have been normalized to the value for hTRF1 (see *SI Appendix*, Materials and Methods for details). Bar heights are given as the mean ± SD from triplicate activity measurements.

To explore the influence of the substrates on DegP oligomerization, we measured z-average diffusion constants (*D*_*z*_) by DLS. We have previously shown *D*_*z*_ values to be sensitive reporters of the higher order oligomers that are formed by DegP in the absence and presence of client proteins such as hTRF1 (28). In this study we have extended our earlier approach, which utilizes a plate reader format DLS instrument, to screen in a high throughput manner for changes to DegP’s oligomeric distribution in response to client binding. This strategy enabled us to investigate multiple client concentrations and solution temperatures in a single experiment and thereby explore a wide range of conditions under which DegP is thought to act as a stress-protective protease and chaperone (5, 11, 33). Initially, *D*_*z*_ values for each of the five substrates in Figure 2A were measured as a function of temperature (5-40 °C) to evaluate their hydrodynamic size and thermal stability (Fig. 2B). As expected, we found a clear correlation between the measured *D*_*z*_ values and the size of the substrate, with smaller substrates diffusing faster than larger ones, according to *D*_*z,hTRF1*_ > *D*_*z,TrpCage-hTRF1*_ > *D*_*z,His-SUMO-hTRF1*_ ≈ *D*_*z,IL6-hTRF1*_ *> D*_*z,MNeon-hTRF1*_ over the temperature range of ∼5-30 °C. Certain substrates (TrpCage-hTRF1, IL6-hTRF1, and MNeon-hTRF1) were found to undergo irreversible unfolding and/or aggregation beyond these temperatures, as evidenced by their DLS autocorrelation functions (*SI Appendix*, Fig. S1). The dramatic shifts to longer autocorrelation function decay times and the appearance of multiphasic behaviour and noisy baselines at these temperatures are consistent with the formation of multiple large scattering species corresponding to high molecular weight protein aggregates (34). The fits of these types of autocorrelation functions are poor (*SI Appendix*, Fig. S1) and give rise to the steeply decreasing and anomalously small extracted *D*_*z*_ values that are shown in Figure 2B over the range of ∼30-40 °C.

We then measured *D*_*z*_ values for proteolytically inactive S210A DegP (to prevent auto and substrate catalysis) in the presence of each substrate as a function of temperature (5-40 °C, Fig. 2C and *SI Appendix*, Fig. S2; colored circles for client-bound data match the client colors in Fig. 2A and Fig. 2B). We selected a 100 μM S210A DegP protomer concentration since this is within the experimentally determined biological range for DegP in the *E. coli* periplasmic space (28) and used three concentrations of each substrate (50, 100, and 200 μM) to assess cage formation at several molar ratios of DegP:substrate (Fig. 2C and *SI Appendix*, Fig S2; note that in Fig. 2C, only the 1:2 monomer:monomer data are shown for clarity). As the substrates are small relative to DegP trimers and cages (∼7 to 33 kDa *vs* ∼140 kDa for a DegP trimer as the smallest DegP oligomer; note that this mass difference increases dramatically upon cage formation) and are not in dramatic molar excess even at the highest employed concentration, the contribution of any unbound client to the measured *D*_*z*_ values is negligible. Therefore the measured *D*_*z*_ values are direct reporters of the formation of client-engaged DegP cages (28). A sample of apo DegP at 100 μM protomer concentration was also included as a reference, highlighting DegP’s oligomeric distribution in the absence of substrate (Fig. 2C, red circles). The measured *D*_*z*_ values for DegP in the absence and presence of the five substrates followed *D*_*z,DegP*_ > *D*_*z,DegP+hTRF1*_ ≈ *D*_*z,DegP+TrpCage-hTRF1*_ > *D*_*z,DegP+His-SUMO-hTRF1*_ > *D*_*z,DegP+IL6-hTRF1*_ > *D*_*z,DegP+MNeon-hTRF1*_ at all temperatures, suggesting the formation of different cage distributions depending on the size of the N-terminal portion of the bound client (Fig. 2C and 2D). Notably, we did not observe changes to the autocorrelation functions for DegP in the presence of the clients over this temperature range that would indicate the formation of non-specific protein aggregates (*SI Appendix*, Fig. S1), as was found for apo TrpCage-hTRF1, IL6-hTRF1, and MNeon-hTRF1, however, the concentrations of free protein required for accurate measurements of *D*_*z*_ values (Fig. 2B, *SI Appendix*, Fig. S1) are much higher than for the complexes in Figure 2C. To rule out the possibility that the chimeric substrates could bind to DegP via regions other than the hTRF1 tag, we prepared constructs containing only the N-terminal portion of the chimeras (*i*.*e*. TrpCage, His-SUMO, IL6, and MNeon) and repeated the DegP binding experiment by DLS (*SI Appendix*, Fig. S3). In all cases the *D*_*z*_ values for S210A DegP in the presence of these constructs did not show any decreases in magnitude and reproduced the shape of the apo S210A DegP reference sample, indicating that cage formation is not induced by these clients without the C-terminal hTRF1 tag. Thus, changes in particle sizes reflect differences in substrate radii and not changes in the mechanism of binding to DegP.

To provide estimates of the cage sizes that correspond to the *D*_*z*_ values measured in the presence of the clients over the experimental temperature range, we calculated temperature dependent diffusion constants for a series of DegP oligomers (3mer, 6mer, 12mer, 24mer, and 60mer, grey dashed lines in Fig. 2C and 2D) based on experimentally determined *r*_*h*_ values for 3mer, 6mer, and 12mer particles, or from the structures of the higher-order cages determined here (see *Cryo-EM structural studies of DegP cages in the presence of engineered substrates*). Consistent with our previous study on DegP oligomerization in the absence of substrate (28), the *D*_*z*_ values for apo DegP under these conditions initially track with those expected for a hexamer at low temperature (∼5-15 °C) and then subsequently decrease as the temperature is elevated, reflecting the higher-order apo oligomers that become populated at higher temperatures (∼25-40 °C). The *D*_*z,DegP+hTRF1*_ data were consistent with the values expected for a 12mer cage, in agreement with our previous report (28). *D*_*z*_ values for DegP in the presence of TrpCage-hTRF1 were slightly below the values for hTRF1, possibly the result of the formation of a small amount of 18mer cages in addition to 12mers (28). For DegP in the presence of the larger substrates (His-SUMO-hTRF1, IL6-hTRF1, and MNeon-hTRF1), we observed fluctuations in the *D*_*z*_ values that were suggestive of substantial redistributions of the DegP cage ensembles in response to increasing temperature (Fig. 2C and *SI Appendix*, Fig. S2). At low temperature (∼5-15 °C), *D*_*z*_ values for DegP bound to His-SUMO-hTRF1 are situated between those expected for 12mer and 24mer cages, consistent with a cage distribution containing a mixture of these (and likely 18mer) particles. At higher temperatures (∼>20 °C), the *D*_*z*,DegP+His-SUMO-hTRF1_ values transition toward and eventually pass slightly below the 24mer values, pointing to the formation of a cage distribution dominated by 24mer particles and a small amount of higher order species. A similar trend was observed for DegP bound to IL6-hTRF1, except that these *D*_*z*_ values are consistent with a largely 24mer cage distribution at lower temperature (∼5-15 °C), shifting toward the values expected for a 60mer cage at high temperature (∼25-40 °C). Finally, the *D*_*z,DegP+MNeon-hTRF1*_ data initially are close to those expected for a 60mer cage at lower temperatures and deflect slightly below the 60mer line pointing to the formation of even higher order particles at high temperatures. This temperature deflection for DegP in the presence of larger substrates may reflect, in part, the unfolding of the hTRF1 subunit of unbound chimeric clients. An increase in the unfolded fraction of the hTRF1 tag would increase the effective binding affinity, as DegP can bind denatured clients preferentially (31), leading to increased saturation of DegP with temperature. Additional copies of clients populating the cages would then force a redistribution of the ensemble toward larger oligomers, ultimately giving rise to the observed deflections in the measured *D*_*z*_ values. Alternatively, formation of larger particles for the His-SUMO-hTRF1, IL6-hTRF1 and MNeon-hTRF1 complexes may simply reflect a temperature dependent re-distribution of cage sizes in a manner analogous to what occurs with apo DegP (28).

Having established that our suite of engineered DegP clients leads to different cage distributions, we next sought to assess how these artificial substrates would be cleaved by protease-active DegP. To obtain a crude estimate of DegP activity toward the clients, we incubated active DegP with each of the substrates at room temperature overnight and analyzed the reaction products using SDS-PAGE (*SI Appendix*, Fig. S4). In all cases, we observed hTRF1 cleavage products exclusively, implying that active DegP digests only the hTRF1 tag. This is most evident for the larger substrates (His-SUMO-hTRF1, IL6-hTRF1, MNeon-hTRF1), where low molecular weight bands corresponding to the hTRF1 tag cleavage products are observed in addition to higher molecular weight bands for the free, intact N-terminal portions of these chimeras. To confirm that these cleavage products are the result of C-terminal hTRF1 tag digestion, we subjected the reaction products to LC-MS analysis. A similar product profile for each of the chimeric substrates was observed, with the accompanying peptide masses matching those expected for cleavage of the hTRF1 tag (*SI Appendix*, Table S2). We also measured activity profiles for each of the substrates using changes in the intrinsic fluorescence of hTRF1 as a reporter. These data revealed similar cleavage rates for hTRF1, His-SUMO-hTRF1, and MNeon-hTRF1, while TrpCage-hTRF1 and IL6-hTRF1 were digested at somewhat slower rates (∼50% and 20% of the hTRF1 rate, Fig. 2E). Differences in activity for the TrpCage-hTRF1 and IL6-hTRF1 constructs support the notion that their N-terminal portions influence DegP protease activity against the bound hTRF1 tag, which could have implications for DegP function in the periplasm (see Discussion).

### Cryo-EM structural studies of DegP cages in the presence of engineered substrates

The correlation between the sizes of the engaged clients and the DegP cages, as evidenced by the DLS-derived *D*_*z*_ values (Fig. 2C and 2D), prompted us to use single-particle cryo-EM to investigate the structures of the substrate-bound assemblies in more detail. To this end we prepared samples of the series of five DegP:client complexes (at 100 μM DegP monomer:200 μM client concentrations to ensure saturation of the cages with substrate) and subjected each to cryo-EM analysis (*SI Appendix*, Materials and Methods). Inspection of the micrographs and 2D class averages for the different complexes examined suggested increasingly large effective DegP cages as a function of increasing client size (*SI Appendix*, Fig. S5), in agreement with our macroscopic analyses of the DegP cage distributions by DLS. For each dataset, particle images were subjected to *Ab-initio* reconstruction and classification (*SI Appendix*, Fig. S5) followed by refinement (35) with appropriate symmetry enforced, depending on the type of cage (*SI Appendix*, Fig. S6, *SI Appendix*, Table S3 and S4, *SI Appendix*, Materials and Methods).

In the presence of the smallest client, hTRF1, a uniform distribution of DegP structures is formed, consistent with tetrahedral 12mer cages (100% of particles, Fig. 3A, *SI Appendix*, Fig S5A), as observed previously (28). When bound to larger substrates (*i*.*e*., the chimeric clients that included the hTRF1 tag), more than one type of DegP cage was visually evident in micrographs (*SI Appendix*, Fig. S5B-D, Fig. 3A), and these were confirmed by *Ab initio* reconstruction (Fig. 3B). For the second smallest client, TrpCage-hTRF1, major (∼92%) and minor (∼8%) populations of 12mer and trigonal bipyramidal 18mer cages were obtained, respectively. For the next largest substrate, His-SUMO-hTRF1, the size distribution was predominantly weighted towards 12mers (∼57%), with a significant shift to a larger fraction of 18mer cages (∼30%), and the appearance of a minor population of octahedral 24mer particles (∼13%). When engaged with the second largest client, IL6-hTRF1, the smallest particles formed were 24mers, a shift in size distribution that likely reflects the volume constraints imposed by encapsulation of the larger IL6 domain of the chimera within the cage interiors. In this case nearly equal populations of two major particle types were obtained, 24mer cages (52%) and a novel pentagonal bipyramidal 30mer class (42%). In addition, a small fraction of the particles (6%) formed an icosahedral 60mer cage that, to our knowledge, has not been observed before either. The particle distribution for DegP bound with the largest client, MNeon-hTRF1, consisted largely of these 60mer cages (77%) and strikingly, also included much larger amorphous DegP assemblies on the order of subcellular organelles in diameter (23% of particles, diameter ∼20-100 nm, *SI Appendix*, Fig. S5D).

**Figure 3.**
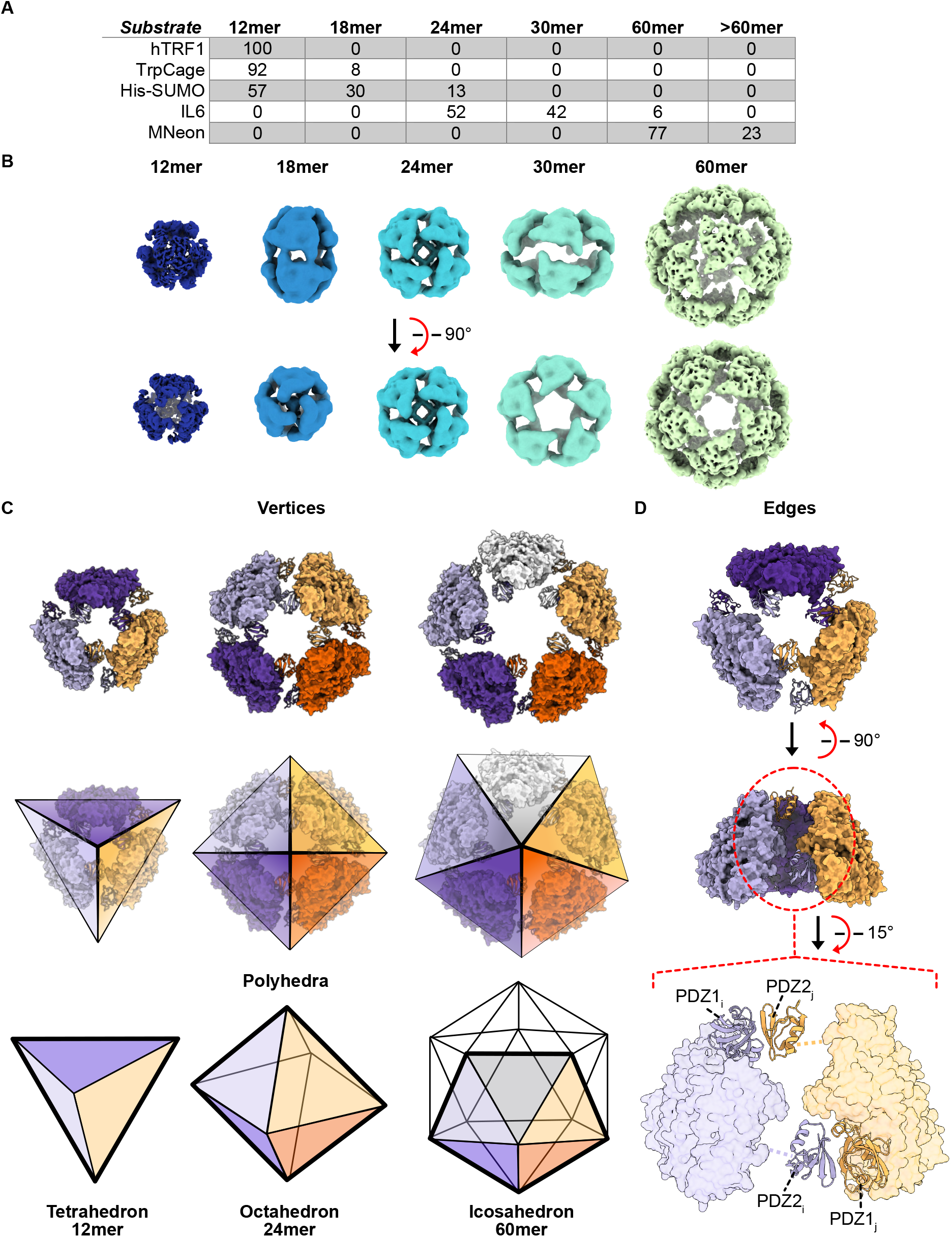
Cryo-EM analysis of the DegP cage structures formed in the presence of chimeric clients. (A) DegP cage particle distributions that are adopted in the presence of each chimeric client, given in percentages of the total number of particles as estimated from *Ab-initio* classification. (B) Cryo-EM maps of the DegP cages adopted in the presence of increasingly large substrates. Maps of a tetrahedral 12mer, a trigonal bipyramidal 18mer, an octahedral 24mer, a pentagonal bipyramidal 30mer, and an icosahedral 60mer are respectively shown from left to right in dark blue to light green. (C, top) DegP cage structures can be approximated as polyhedra by considering the trimer subunits as triangles. Views along the respective 3-, 4-, and 5-fold symmetry axes of the 12mer, 24mer, and 60mer DegP cages from (B) (top and middle; rear subunits are not shown for clarity and representative polyhedral shapes are overlaid in the middle row) highlight that the cage polyhedra (bottom, bold outlines match the edges of the views in the middle row) are assembled via edge-wise interactions between trimers. (D) The DegP cage structures are formed by inter-trimer PDZ1^*i*^:PDZ2^*j*^ domain interactions. To illustrate, the formation of a single edge connecting light purple and orange trimers in the 12mer (top and middle) occurs through a pair of PDZ1^*i*^:PDZ2^*j*^ interactions (bottom).

Cryo-EM maps of the 12mer, 18mer, 24mer, 30mer, and 60mer DegP complexes are highlighted in Figure 3B. Notably, for the extremely large and irregular assemblies identified in the micrographs of the DegP:MNeon-hTRF1 sample (>60mer, *SI Appendix*, Fig. S5D), we were not able to obtain particle classes with sufficient populations for reconstruction. In all other cases structures could be obtained, and they establish that cages are formed through inter-trimer PDZ1^*i*^:PDZ2^*j*^ domain interactions, with the protease domain catalytic cores pointing towards the cage interiors (Fig. 3B and 3C), as was found previously in other studies of substrate-engaged DegP assemblies (18, 25, 27). In none of the structures were we able to clearly identify DegP’s protease domain LA loops (residues 36-81) or the PDZ1-PDZ2 linker sequences (residues 359-374), both of which are known to be highly plastic (25, 36). In addition, there was no obvious density for the N-terminal portions of the chimeric substrates, probably also as a result of their high levels of conformational flexibility and the lower overall resolution of most of the obtained maps in the presence of these clients. We wish to point out that in Figure 3B, although certain cages appear to contain density within their interiors (*e*.*g*. the 24mer), this is the result of the symmetry enforcement during refinement and does not correspond to encapsulated substrates or DegP’s protease domain loops. An exception was for the 12mer cage structure derived from the DegP:hTRF1 sample, where regions of the bound hTRF1 chains could clearly be identified (see *hTRF1 forms interactions that stabilize the DegP 12mer cage and promote catalysis*).

As illustrated in Figure 3C, each of the DegP structures can be pictured as a polyhedron with triangular faces that correspond to the constituent trimer building blocks. The edges of the polyhedra are formed by inter-trimer PDZ1^*i*^:PDZ2^*j*^ domain interactions (Fig. 3D top and middle, with edge interactions indicated for the light purple and light orange trimer structures), where the PDZ domain pairings occur at the top and bottom of each edge (Fig. 3D bottom showing pairs of ribbon PDZ domains). Different cage sizes, and thus surface curvatures, are created by exploiting these edgewise PDZ domain interactions and the flexible protease-PDZ1 and PDZ1-PDZ2 domain linkers within a protomer that enable an expansion of the gaps in the vertices of the polyhedra (Fig. 3C top). In order to better visualize the exquisite structural plasticity of DegP’s linker sequences and PDZ1^*i*^:PDZ2^*j*^ domain interactions, 3D variability analysis (37) was performed for the 12mer and 60mer structures in which continuous protein motions are described as linear additions of density to a consensus map. These cages were chosen so as to explore the range of motions that complexes of drastically different sizes might experience. The major components of structural variability derived from the analysis revealed that the 12mer and 60mer cages undergo extensive twisting and flexing movements respectively (*SI Appendix*, Supplementary Movie 1 and 2). These motions, arising from the flexible trimer building blocks, could be important for DegP cage assembly and client processing (see Discussion).

### hTRF1 forms interactions that stabilize the DegP 12mer cage and promote catalysis

With the establishment of the general structural features of the DegP cages that are formed using the client proteins discussed above (*Cryo-EM structural studies of DegP cages in the presence of engineered substrates*), we next asked whether specific information could be obtained with reference to DegP:substrate interactions from our cryo-EM data. We focused on the DegP:hTRF1 dataset as it had the most uniform particle distribution and was of the highest overall quality. The trimer map was generated by symmetry expansion of the 12mer map followed by local refinement with a mask over a single trimer, which increased the global resolution from 3.1 to 2.6Å (Fig 3B and Fig 4A, *SI Appendix*, Fig. S6A). The increase in resolution from the local refinement implies that there is flexibility between different trimer faces, consistent with the 3D variability analysis that was performed (*SI Appendix*, Supplementary Movie 1 and 2). Importantly, at the level of detail afforded by the local refinement, we could clearly identify a number of novel structural interactions between DegP protomers and hTRF1 chains (Fig 4B-D). Well-defined density is observed in the cryo-EM map for the C-terminal halves of the hTRF1 chains (27 residues, Fig. 4B; the segments of the hTRF1 chains that could be visualized are shown in pink ribbon, the structure of natively folded hTRF1 (32) is shown to the bottom right as a reference with native helices numbered and colored according to the sequence bound to DegP). As mentioned, density was not observed for the N-terminal portions of the hTRF1 chains, presumably due to their flexibility within the cage interiors. The C-terminal halves of hTRF1 clients are arranged in a propeller-shape, as shown for the trimer in Figure 4B, where each hTRF1 molecule is bound in a bi-partite manner to the PDZ1 domain of a given protomer and the protease domain of an adjacent protomer (counterclockwise in Fig. 4B) within the same trimer. This type of binding mode for DegP clients has previously been proposed based on partial electron density in X-ray studies of peptides engaged to 12mer and 24mer DegP particles (24, 25).

**Figure 4.**
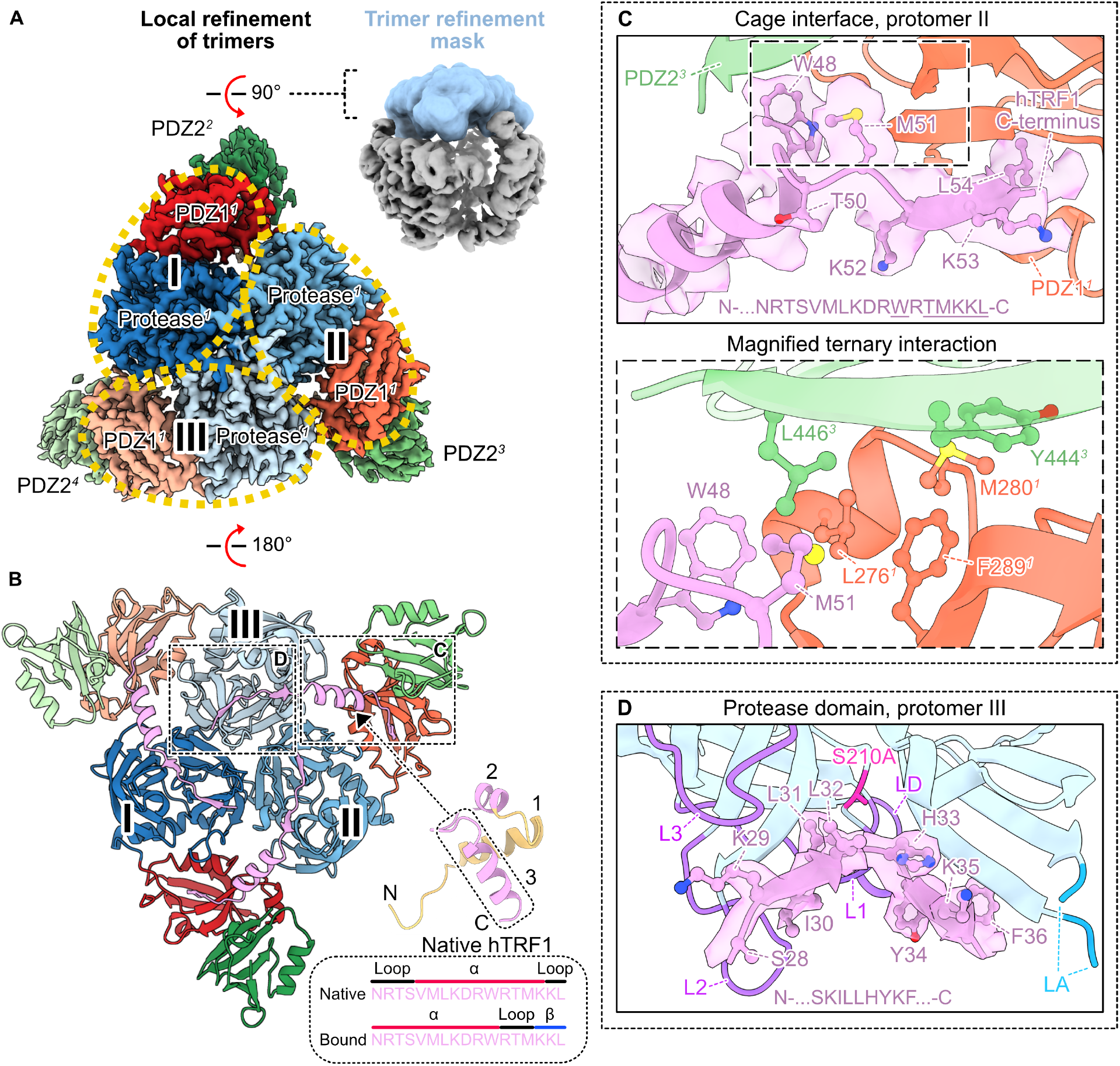
The structure of a 12mer DegP cage reveals interactions between DegP and hTRF1. (A) Cryo-EM density maps for an hTRF1-bound DegP 12mer. Local refinement of a single trimeric face of the 12mer (the refinement mask is indicated in blue in the top right) improved the resolution to 2.6Å (left). The protease and PDZ1 domains in each of the three protomers (protomers are indicated as I, II, and III) of the displayed trimer (denoted as trimer 1 by the numerical superscript on the domain labels; that is, all 9 domains of the 3 protomers comprising trimer I are denoted by 1, those of trimer II by 2, and so forth) are outlined with gold ovals and the protomer numbers are indicated in bold. The PDZ2 domains from the protomers within trimer 1 are not visible in this orientation; they form contacts establishing other oligomeric interfaces in the 12mer. PDZ2^*j*^ domains contributed by trimers 2-4 can be seen along the outside of trimer 1 and are denoted by PDZ2^2-4^ to indicate that they derive from trimers distinct from trimer 1. (B) Structural model of hTRF1 bound to a DegP trimer within the 12mer cage. Counter-clockwise from protomer I, the protease and PDZ1 domains of each protomer in the DegP trimer are shown in dark to light blue (protease domains of successive protomers within the trimer) and red to light orange (successive PDZ1 domains). The PDZ1 domains form PDZ1^*i*^:PDZ2^*j*^ contacts with PDZ2 domains (shown in dark to light green) contributed by protomers from other trimers. The C-terminal portion of hTRF1 that interacts with DegP is shown in pink. For reference, the natively folded hTRF1 structure (32) is shown in the bottom right with the helices numbered and structure colored according to the sequence that we identified as bound to DegP (pink for observed, yellow for not observed). Secondary structure diagrams for residues surrounding and including the third helix in native hTRF1 are indicated to highlight the remodeling of this helix upon binding to DegP. (C, top) Closeup of the cage interface formed by hTRF1 and PDZ1^*1*^ (protomer II in panel (B)) and PDZ2^*3*^ domains. The map for the hTRF1 chain at this interface (corresponding to the amino acid sequence indicated at the bottom of the panel) is shown overlaid with the structural model. Residues with well-defined sidechain density have been modelled and are shown as balls and sticks; these are denoted by the underlined portion of the hTRF1 sequence (N-…<u>W</u>R<u>TMKKL</u>-C). Colored balls correspond to sidechain heteroatoms (nitrogen in blue, sulfur in yellow, and oxygen in red). (C, bottom) Expanded region of (C, top) detailing the side chains of hTRF1 and PDZ1^*1*^ and PDZ2^*3*^ domains that contribute to the cage interface. (D) Closeup of the interactions between hTRF1 and the protease domain of protomer III from trimer 1 in (B). The hTRF1 sequence for which we observed density is indicated at the bottom of the panel. The catalytic serine is indicated by the red stick (The inactive S210A mutation is employed here). The protease domain L1, L2, L3, and LD loops that have undergone a conformational change upon binding substrate to achieve the catalytically active conformation of DegP are shown in purple. The base of the long LA loop for which we did not observe density is indicated in cyan.

Interactions between an hTRF1 client molecule and the individual domains of DegP trimers in the context of the 12mer structure are highlighted in Figures 4C and 4D. The hTRF1 chains are engaged at their C-termini to the peptide-binding clefts of the PDZ1 domains (Fig. 4B, with the three PDZ1^*1*^ domains from trimer 1 shown as red, orange, and light orange ribbon). Docking of each hTRF1 C-terminus (N-…KKL-C, Fig. 4C) is, in part, facilitated by the formation of a non-native β-strand between the C-terminal three residues of hTRF1 and an existing *β*-strand of the PDZ1 domain whose edge faces the binding groove. These hTRF1 residues that extend the PDZ1 strand comprise the final turn of the client’s natively folded 3^rd^ α-helix that is unwrapped upon binding (Fig. 4B bottom right, see the secondary structure diagrams for the sequence corresponding to the hTRF1 structure outlined in the dashed box). In addition, the client complex is stabilized through the insertion of hTRF1’s terminal Leu sidechain and carboxylate group into the peptide-binding pocket. Remarkably, for the segment of the hTRF1 chain that bridges the PDZ1 and protease domain binding sites (*i*.*e*. N-…NRTSVMLKDRW…-C), we observed a well-defined helix formed from the majority of the third α-helix in native hTRF1 and several preceding residues from a loop that adopt an additional helix turn when bound to DegP (Fig. 4B bottom right, see comparison of native and bound secondary structure diagrams for this segment).

Remodeling of this native hTRF1 helix upon binding to DegP, effectively shifting its register, appears to be important for stabilizing PDZ1^*i*^:PDZ2^*j*^ domain interactions between DegP trimers by enabling the formation of a ternary hTRF1:PDZ1^*i*^:PDZ2^*j*^ cage interface (Fig. 4C). For example, Met51, which in natively folded hTRF1 resides in the final helix (Fig. 4B), is restructured so that its sidechain contributes to the hydrophobic cage interface formed, as do key residues in the PDZ1^*i*^ (Leu276^*i*^, Met280^*i*^, and Phe289^*i*^) and PDZ2^*j*^ domains (Tyr444^*j*^ and Leu446^*j*^; PDZ2^*j*^ domains are shown as dark green, green, and light green ribbon). The sidechain of nearby Trp48, which remains in its native helical conformation in the bound form (Fig. 4B), also contributes to the packing of this hydrophobic ternary cluster (Fig. 4C). Thus, the active restructuring of the client through engagement by DegP directly contributes to the stability of the complex. These ternary hTRF1:PDZ1^*i*^:PDZ2^*j*^ interactions also stabilize the packing of PDZ1^*i*^:protease^*i*^ domains (Fig. 4B), forming a conformation that has been shown to promote the activation of DegP towards bound peptides (24).

The hTRF1 chain makes a substantial number of additional contacts with the protease domain of a neighbouring protomer in the same trimer (Fig. 4B and 4D, protease domains are shown as dark blue, blue, and light blue ribbon). Notably, regions of an hTRF1 chain bound to the protease domain (N-…SKILLHYKF…-C) form a V-shape through the adoption of two non-native β-strands that extend a pair of β-sheets proximal to the catalytic center of DegP (Fig. 4D). This interaction arises through the disruption of native hTRF1’s 2^nd^ α-helix (Fig. 4B bottom right), and places the hTRF1 chain in a bent conformation with DegP’s active site catalytic serine (in this case alanine, red stick in Fig. 4D) positioned for efficient proteolysis at the base of the V-shape formed by the hTRF1 peptide backbone. The hTRF1 cleavage products derived from proteolysis by active DegP (*SI Appendix*, Table S2) correlate with the engagement of the hTRF1 chain in this manner. In addition to the well-defined bound-state structure that we identified for the hTRF1 chains, DegP’s protease domains were found to be in their catalytically active conformations. This is evidenced by the positions of the L1, L2, L3, and LD loops (Fig. 4D purple loops, note the base of the unobserved LA loop is colored cyan) surrounding the active sites which permits the alignment of the catalytic centers for client proteolysis (18).

## Discussion

DegP functions as both a protease and a chaperone (5, 10, 18, 38), playing important housekeeping and virulence-promoting roles in the periplasm of gram-negative bacterial pathogens. DegP, thus, represents an important antibiotic target (23), and a recent study has shown that its overactivation by small molecules that mimic the C-terminus of a trimeric OMP client is a viable route for inhibiting bacterial growth (39). Our structural study of client-engaged DegP oligomers presented here suggests that ternary client:PDZ1^*i*^:PDZ2^*j*^ interfaces could be targeted by compounds that augment DegP cage stability to promote enhanced proteolysis, potentially leading to bacterial cell death. Collectively, the many forms of DegP cages that we and others have identified (6, 18, 25–28) offer a rich structural landscape for potential exploitation in the development of antibacterial therapies.

In this study we have used a combined DLS/cryo-EM approach to explore the types of cages adopted by DegP in the presence of a series of client proteins with N-terminal domains of increasing hydrodynamic radii. Notably, these techniques have allowed the simultaneous and facile identification of multiple oligomeric species which form in response to the engaged cargo, leading to the identification of a pair of novel structural ensembles. These include pentagonal bipyramidal 30mer and icosahedral 60mer cages, the latter similar to a viral capsid (40). Particle images were also obtained for DegP assemblies which are on the order of subcellular organelles in diameter (∼100 nm), although structures could not be elucidated in this case. Our results establish that DegP can adopt a variety of cage configurations whose sizes depend on the size of the bound substrate, and that large biologically relevant substrates can be accommodated. For example, the 60mer and amorphous higher-order DegP assemblies could play roles in the transport of very large cargo within the periplasm, notably virulence factors such as autotransporter proteins whose molecular weights are on the order of 100 kDa (15, 41). Due to their large folded structures, it is thought that autotransporters are kept in a partially unfolded state within the periplasm to facilitate their secretion to the cell exterior (42, 43). The enormous DegP assemblies observed here could serve as protective vesicles that stabilize the denatured state of the autotransporter chain for subsequent secretion.

In a previous study of the structural dynamics of DegP we showed that an ensemble of complexes was present in solution in the absence of substrate, and that this ensemble was likely important to facilitate a timely response to cell stresses that would involve the interaction of the protease-chaperone with a wide range of clients (28). In the present study we have highlighted the importance of dynamics in the context of substrate-engaged DegP complexes as well. Notably, an ensemble of different particle types, which can be most simply described in terms of polyhedra, are found for many of the complexes, with the distribution of particle sizes changing in response to the substrate. The variety of structures that can be achieved with the same trimer building blocks, giving rise to the observed wide variability in particle architectures, derives from the inherent flexibility of the connections between the protease and PDZ domains of a given protomer. Moreover, even in the context of a single particle type, for example the tetrahedral 12mer or icosahedral 60mer DegP structures formed with hTRF1 and MNeon-hTRF1 respectively, there are extensive twisting and breathing motions, as established through 3D variability analyses that, again, derive from the inherent plasticity of the protomer and trimer units (*SI Appendix*, Supplementary Movie 1 and 2). This structural variability may be important for reorganization of cage intermediates in the assembly pathway and might also permit the reorientation of bound clients once inside DegP for generating proteolytically competent conformations. These motions could additionally facilitate client entry and the subsequent egress of cleaved polypeptides. Alternatively, the inherent flexibility of the cages may lead to nuanced structural changes as a function of the number of bound substrates, potentially regulating the function of DegP as either a chaperone or protease.

Compared to crystallographic studies of DegP assemblies that only observed client density separately in the protease and PDZ1 binding sites (18, 24, 25), our structural studies of the 12mer DegP complex in the presence of hTRF1 allowed observation of connectivity both within and between sites on hTRF1 involved in binding, showing that, at least in this case, the bound substrate is not completely unfolded. Indeed, hTRF1’s C-terminal half is remodeled to play an active role in stabilizing the structure of the complex through interactions at the PDZ1^*i*^:PDZ2^*j*^ interfaces where the three C-terminal residues of hTRF1 extend an existing PDZ1 *β*-sheet at the binding site, leading to unwinding of a single turn of a helix in the native hTRF1 structure and subsequent addition of a turn at the opposite helix end. Additionally, catalysis is facilitated through the formation of a V-shaped pose of the hTRF1 chain that is stabilized through the adoption of two non-native hTRF1 β-strands that extend a pair of β-sheets near the catalytic center of DegP. Interestingly, our LC-MS analyses of the hTRF1 cleavage products derived from proteolysis by active DegP (*SI Appendix*, Table S2) indicate that hTRF1 can bind to DegP in more than one register, underscoring a possible functional interplay between bound client dynamics and catalysis.

This work paints a picture of a highly dynamic DegP – substrate system, with structural flexibility at the level of both receptor and ligand components. DegP assemblies can be thought of as highly dynamic and adaptive cages that are governed by a complex, client-dependent free energy landscape, where cage assemblies redistribute according to the number of bound substrate copies, client sizes, and solution conditions such as temperature. *In vivo*, DegP cage distributions are most likely constantly in flux within the bacterial periplasm in response to oscillations in misfolded client levels that are the result of a variety of cellular stressors including heat (5), oxidative (11), and osmotic shock perturbations (12). An understanding of the complexities of DegP’s free energy landscape is an important first step in the design of molecules to regulate its function and, potentially, to mitigate the virulence of classes of bacterial pathogens.

## Acknowledgements

This research was funded by a grant from the Canadian Institutes of Health Research, FDN-503573 (L.E.K.). Dr. John Rubinstein (Hospital for Sick Children, Toronto) is thanked for facilitating the use of a Tecnai F20 microscope and computing resources. R.W.H and J.D.T. are grateful to CIHR and the Hospital for Sick Children Research Institute for post-doctoral fellowships.

## Materials and Methods

Details of protein expression and purification, and all experiments, along with data analysis and fitting, are provided in *SI Appendix*.

## Data Availability

All relevant data, outside of structural coordinates, are included in the paper and in *SI Appendix*.

## Data deposition

Electron microscopy maps and atomic models have been deposited to the Electron Microscopy Data Bank and the Protein Data Bank under the accession nos. and PDB ID codes EMD-XXXX, XXXX (12mer); EMD-XXXX, XXXX (3mer local refinement); EMD-XXXX, XXXX (18mer); EMD-XXXX, XXXX (24mer); EMD-XXXX, XXXX (30mer); EMD-XXXX, XXXX (60mer).

