## Supplementary Information for "Flexible client-dependent cages in the assembly landscape of the periplasmic protease-chaperone DegP"

### Materials and methods

*Plasmid constructs and cloning*

The genes for expression of mature DegP (residues 27-474 lacking the periplasmic signal sequence and renumbered here as 1-448) and the DNA binding domain of human TRF1 (residues 378-430, renumbered here as 1-54 including an additional N-terminal Gly residue; see Table S1 for the construct sequence) were obtained as described previously (1). The gene sequences for the chimeric clients TrpCage-hTRF1, IL6-hTRF1, and MNeon-hTRF1 were obtained by attaching the nucleotide sequence for hTRF1 to the 3' ends of the TrpCage, IL6, and MNeon genes respectively (see Table S1 for the protein sequences derived from these genes). We note that the His-SUMO-hTRF1 gene was not prepared in this manner as it could be produced by subcloning the sequence for hTRF1 into a plasmid backbone containing an N-terminal $His_6$-SUMO tag, as outlined below. For TrpCage-hTRF1 and His-SUMO-hTRF1, a linker sequence corresponding to a single Gly residue was included between the N-terminal protein and the C-terminal hTRF1 tag to allow for conformational freedom between the two domains of these chimeric clients. The genes for the chimeric constructs were synthesized as codon-optimized gBlocks® (Integrated DNA Technologies) containing 5' and 3' overhangs for subcloning. All expression constructs were prepared by subcloning the genes of interest using the Gibson Assembly® procedure (New England BioLabs Inc.) into kanamycin resistance pET plasmids with N-terminal $His_6$ affinity tags for purification. For MNeon-hTRF1, a pET29 plasmid was used with the N-terminal $His_6$ tag containing a C-terminal ENLYFQG motif that enabled cleavage of the affinity tag by TEV protease. All other constructs were subcloned into a *Champion* pET SUMO plasmid (hTRF1, TrpCage-hTRF1, IL6-hTRF1) which contains a sequence corresponding to an N-terminal $His_6$-SUMO tag. The expression constructs for IL6 and MNeon were generated by mutating the first hTRF1 residue in the IL6-hTRF1 and MNeon-hTRF1 plasmids to a stop codon using the QuikChange site-directed mutagenesis method. Isolated TrpCage was not produced in this way; instead, it was obtained from GenScript as a synthetic peptide. It was not necessary to introduce a stop codon into the His-SUMO-hTRF1 construct to obtain a plasmid for expression of His-SUMO alone as this protein could be obtained during the purification of His-SUMO-hTRF1 as described in the following section.

*Protein expression and purification*

All proteins were expressed by growing transformed cells to an $OD_{600}$ of ~0.6-1.0 at 37 °C in LB media at which point IPTG was added to 0.2 mM. Protein expression proceeded for a further ~16-20 hours at 23 °C, except for protease active DegP which was expressed for only 12 hours at 16 °C to mitigate autocatalysis. Other than DegP and IL6, all proteins were expressed in *Codon+ E. coli BL21*(*DE3*) cells. As DegP and IL6 contain native disulfide bonds (1, 2), these proteins were expressed in *SHuffle T7 Competent E. coli cells* which produce disulfide isomerases that promote disulfide bond formation during cytoplasmic expression.

Protein purification proceeded in general according to (1) cell lysis and centrifugation, (2) nickel affinity chromatography, (3) $His_6$-SUMO tag cleavage, (4) an intermediate column chromatography step, and (5) size exclusion chromatography. For protease active DegP, all steps post-refolding from denaturing conditions (see below) were performed on ice or at 4 °C to diminish autocleavage. Cells were lysed on ice in lysis buffer (for most proteins this consisted of 100 mM $NaH_2PO_4$, 500 mM NaCl, pH 8.0; see below for exceptions) and subjected to centrifugation at 9000 rpm for 15 minutes to clear the lysate. Filtered supernatant was loaded onto a nickel affinity purification column equilibrated with lysis buffer. The column was washed with buffer containing

10-30 mM imidazole pH 8.0 (3 rounds of 50-100 mL) to remove contaminant proteins. $His_6$-SUMO-tagged proteins of interest were eluted with lysis buffer containing 500 mM imidazole. For DegP, hTRF1, and TrpCage-hTRF1, the cell lysis and nickel chromatography steps were performed under denaturing conditions where 6M Gdn-HCl was included in the lysis buffer. This was implemented to remove bound substrates from DegP (1) and to protect hTRF1 and TrpCage-hTRF1 from protease cleavage in the earlier stages of the purification. DegP was eluted under denaturing conditions, while hTRF1 and TrpCage-hTRF1 were refolded on-column using a single five column volume wash step to 0M Gdn-HCl prior to elution with imidazole. DegP was subsequently refolded according to our previously published protocol (1). $His_6$-SUMO tags were cleaved by incubation of the (refolded) elution fractions with Ulp1 protease for ~1 hour, except for His-SUMO-hTRF1 where the tag was left attached. Hydrophobic interaction chromatography was used as an intermediate purification step for DegP, IL6, and MNeon using a 5 mL Butyl HiTrap column according to our published method (1). Briefly, nickel eluates were loaded onto the column after addition of solid ammonium sulfate to 1M. A stepped gradient from 1M to 0M ammonium sulfate in a base buffer of 25 mM $NaH_2PO_4$, 1 mM EDTA, pH 7.0 was used to elute the proteins of interest. Cation exchange chromatography was performed as an intermediate purification step for hTRF1, TrpCage-hTRF1, His-SUMO-hTRF1, IL6-hTRF1, and MNeon-hTRF1 using a 5 mL SP Sepharose column. The nickel eluates containing these proteins were diluted with buffer (25 mM $NaH_2PO_4$, 1 mM EDTA, pH 7.0) until the total salt concentrations were estimated to be lower than ~50-100 mM to ensure binding to the column. A gradient from 0 to 1M NaCl over five column volumes in a background of 25 mM $NaH_2PO_4$, 1 mM EDTA, pH 7.0 led to the elution of the protein of interest. Notably, for the proteins whose $His_6$-SUMO tags were cleaved in a prior step by Ulp1 protease, this intermediate chromatography step permitted collection of the freed $His_6$-SUMO tags in the flow-through fraction, which could be independently gel-filtrated for use in the DLS control experiments (*SI Appendix*, Fig. S3). Elution fractions from the intermediate chromatography step were concentrated and subjected to size-exclusion chromatography in 25 mM $NaH_2PO_4$, 200 mM NaCl, 1 mM EDTA, pH 7.0 for final purification (30 mL format Superdex 75 and 200 Increase columns were used for client proteins and DegP respectively). In all cases, the purified proteins were buffer exchanged into the experimental buffer consisting of 25 mM $NaH_2PO_4$, 200 mM NaCl, 1 mM EDTA, pH 7.0 via Amicon Ultra concentrators. The concentrations of purified samples were calculated from their absorbances measured using a NanoDrop spectrophotometer (ThermoFisher) and their extinction coefficients at 280 nm (24 980, 31 970, 26 470, 35 200, and 69 330 $M^{-1}$ $cm^{-1}$ for hTRF1, TrpCage-hTRF1, His-SUMO-hTRF1, IL6-hTRF1, and MNeon-hTRF1 respectively) obtained from the ExPASY protein parameters calculator (https://web.expasy.org/protparam/protparam-doc.html). In order to reduce autocatalysis prior to use in activity assays, protease active DegP was lightly concentrated to 50 μM, aliquoted, flash frozen, and stored at -80 °C.

*DLS measurements and autocorrelation analysis*

DLS autocorrelation functions were measured using a plate reader format Wyatt DynaPro DLS instrument and analyzed according to the cumulants method to extract z-average diffusion constants ($D_z$ values) as previously described (1, 3, 4). Briefly, samples were prepared by dilution of concentrated protein stocks in a buffer of 25 mM $NaH_2PO_4$, 200 mM NaCl, 1 mM EDTA, pH 7.0. To remove contaminating dust, the experimental buffer was filtered (0.2 μm) and the sample tubes and the DLS plate wells were evacuated with compressed air prior to use. Prior to loading on the DLS plate, samples were centrifuged at 17500 rpm for 15 minutes to pellet any large aggregates or dust. Wells containing 20 μL of each sample were capped with 10 μL infrared spectroscopy-grade paraffin oil to inhibit sample evaporation during the temperature ramp experiments. Experiments utilized S210A DegP at a protomer concentration of 100 μM and client concentrations of 50, 100, and 200 μM. Autocorrelation functions (averaged from 25 individual transients) for each sample well were recorded from 5-50 °C in 2.5 °C increments. The average autocorrelation functions were numerically fitted using in-house Python scripts to extract $D_z$ values (1).

*Protease activity assays*

Crude activity assays were performed in 25 mM $NaH_2PO_4$, 200 mM NaCl, 1 mM EDTA, pH 7.0 by incubating 100 μM of the client proteins at 25 °C overnight in the absence and presence of 10 μM DegP. The reaction products were resolved by SDS-PAGE and stained with Coomassie Brilliant Blue for visualization. The gel image in *SI Appendix*, Figure S4 was taken with a Bio-Rad GelDoc EZ imager. The reaction products were also subjected to liquid chromatography-mass spectrometry (LC-MS) analysis at the University of Toronto Department of Chemistry AIMS laboratory in order to identify cleavage fragments of hTRF1 derived from proteolysis by active DegP. The resultant masses corresponding to representative hTRF1 tag digestion products are indicated in *SI Appendix*, Table S2.

Protease activity assays were additionally performed using changes in the intrinsic fluorescence of the hTRF1 tags of the clients as a reporter for cleavage by DegP. A Synergy Neo II plate reader (BioTek) with an excitation wavelength of 295 nm (slit width of 5 nm) and an emission wavelength of 335 nm (slit width of 10 nm) was used to measure the cleavage of the client proteins at 23 °C. Reactions were performed with 2.5 μM DegP and 25 μM substrate protein in 100 μL volumes. Data were recorded at 1 minute intervals for a total of 50 minutes. Initial reaction velocities were obtained as the fitted slopes of the linear portions of the data corresponding to the first ~10-20 minutes of the experiment, depending on the substrate, and were corrected for photobleaching of the hTRF1 tag using an independent measurement of hTRF1 alone. Velocities shown in Figure 2E were normalized to the rate of cleavage of hTRF1 by DegP.

*Single particle electron cryo-microscopy*

*(i) Preparation of electron cryo-microscopy samples*

To prepare proteins for imaging using cryo-EM, nanofabricated holey gold grids were glow discharged in air for 15 seconds, the samples were applied to the grids, and then the grids were blotted for between 15-20 seconds in a modified mark III FEI Vitrobot at 4 °C and ~100% humidity before plunge freezing in a 60:40 propane:ethane mixture held at liquid nitrogen temperature. For each sample, 2.5uL of protein in DLS buffer was applied to holey gold grids with a hole size of ~2 um. For all samples the S210A DegP concentration was 100 μM, and each client protein (hTRF1, TrpCage-hTRF1, His-SUMO-hTRF1, IL6-hTRF1, and MNeon-hTRF1) was added in excess at 200 μM concentration.

*(ii) Electron microscopy*

Datasets for S210A DegP in the presence of TrpCage-hTRF1, His-SUMO-hTRF1, and IL6-hTRF1 were collected on a FEI Tecnai F20 electron Microscope operating at 200 kV and equipped with a Gatan K2 summit direct detector device used in electron counting mode at 400 frames/sec. Movies were recorded as 30 fractions over a 15 second exposure. Defocuses ranged from ~1 to 3 μm. Movies were recorded at a nominal magnification of 25,000× corresponding to a calibrated pixel size of 1.45 Å and with an exposure rate of ~5 electrons/pixel/s, and a total exposure of ~35 electrons/Å$^2$. A total of 166 (TrpCage-hTRF1), 119 (His-SUMO-hTRF1), and 189 (IL6-hTRF1) movies were collected using the Digital Micrograph software package.

For S210A DegP in the presence of hTRF1 and hTRF1-MNeon, datasets were collected on a Titan Krios G3 electron microscope (Thermo Fisher Scientific) operated at 300 kV and equipped with a prototype Falcon 4 direct detector device camera. Movies were recorded at a nominal magnification of 75,000× corresponding to a calibrated pixel size of 1.03 Å and with an exposure rate of ~5.5 electrons/pixel/s, and a total exposure of ~45 electrons/Å$^2$. 4,383 (hTRF1) and 1,185 (MNeon-hTRF1) movies were collected using the automated EPU software package. Movies were recorded as 29 exposure fractions over an 8.7 second exposure.

*(iii) Image analysis*

All image analysis was carried out in *cryoSPARC* v3 (5). For all datasets patch-based alignment and exposure weighting was carried out and the resulting averages of frames were used for patch-based contrast transfer function (CTF) determination and particle picking. Templates for automated particle selection were generated by 2D classification of manually selected particles. The particle images obtained for S210A DegP in complex with hTRF1, TrpCage-hTRF1, His-SUMO-hTRF1, IL6-hTRF1, and MNeon-hTRF1 were extracted in 256x256, 256x256, 256x256, 360x360, and 512x512-pixel box sizes, respectively. Particle images for the His-SUMO-hTRF1, IL6-hTRF1, and MNeon-hTRF1 samples were Fourier cropped to 160x160, 180x180 and 256x256-pixel box sizes, respectively. For all datasets, particles images were subject to *Ab initio* reconstruction and classification as well as 2D classification. The population estimates from the *Ab initio* reconstruction and classification were used to generate the table in Figure 3A. For the hTRF1 dataset, the particle images from the *Ab initio* class corresponding to the 12mer (483,190 particle images) were used in a refinement with enforced tetrahedral symmetry leading to a global resolution of 3.1Å. After refinement, we performed symmetry expansion followed by local refinement with a mask over a portion of the molecule corresponding to a trimer, increasing the resolution to 2.6Å. Since the DegP:TrpCage-hTRF1 sample consisted mainly of 12mer as well, and because we were able to obtain a high resolution 12mer structure from the DegP:hTRF1 sample, no further analysis beyond particle population estimates was performed on the DegP:TrpCage-hTRF1 dataset. For the His-SUMO-hTRF1 dataset, *Ab initio* classes corresponding to the 18mer and 24mer (4,136 and 1,775 particle images) were used in refinements with enforced $D_3$ and octahedral symmetry respectively to generate maps with ~12 and ~14Å resolution. For the IL6-hTRF1 and hTRF1-MNeon datasets, *Ab initio* classes corresponding to the 30mer and 60mer (5,592 and 14,443 particle images) were used in refinements with enforced $D_5$ and icosahedral symmetry respectively, yielding maps with ~14 and ~10Å resolution. 3D variability analyses (6) were performed on the 12mer and 60mer particle datasets (produced from the hTRF1- and MNeon-hTRF1-enaged DegP samples, respectively) to explore cage motions. This analysis was carried out using particle images from refinements where tetrahedral (12mer) and icosahedral (60mer) symmetry were enforced. Twisting and breathing motions of the cages were revealed and examples of these are shown in *SI Appendix*, Supplementary Movies 1 and 2.

*(iv) Model Building*

An atomic model for the client-engaged asymmetric unit of DegP consisting of a protease$^i$ domain, an hTRF1-bound PDZ1$^i$ domain, and an associated PDZ2$^j$ domain (*i* and *j* refer to protomers from separate trimers) was first built in *Coot* (7) using the 2.6Å map of the locally-refined hTRF1-bound DegP trimer. The asymmetric unit obtained from the crystal structure of a peptide-bound DegP 12mer (PDB 6JJO) (8) was used as a starting point for the model. The optimal asymmetric unit from *Coot* was then applied in a *Rosetta* (9) refinement where $C_3$ symmetry was imposed to generate a structural model of the hTRF1-bound DegP trimer. Statistics from this refinement are reported in *SI Appendix*, Table S4. The 12mer, 18mer, 24mer, 30mer, and 60mer cage models were generated by propagating the refined hTRF1-bound trimer using the appropriate symmetry operations in *UCSF Chimera* (10). Figures and movies of maps and models were generated in *UCSF ChimeraX* (11).

### Supplementary Figures

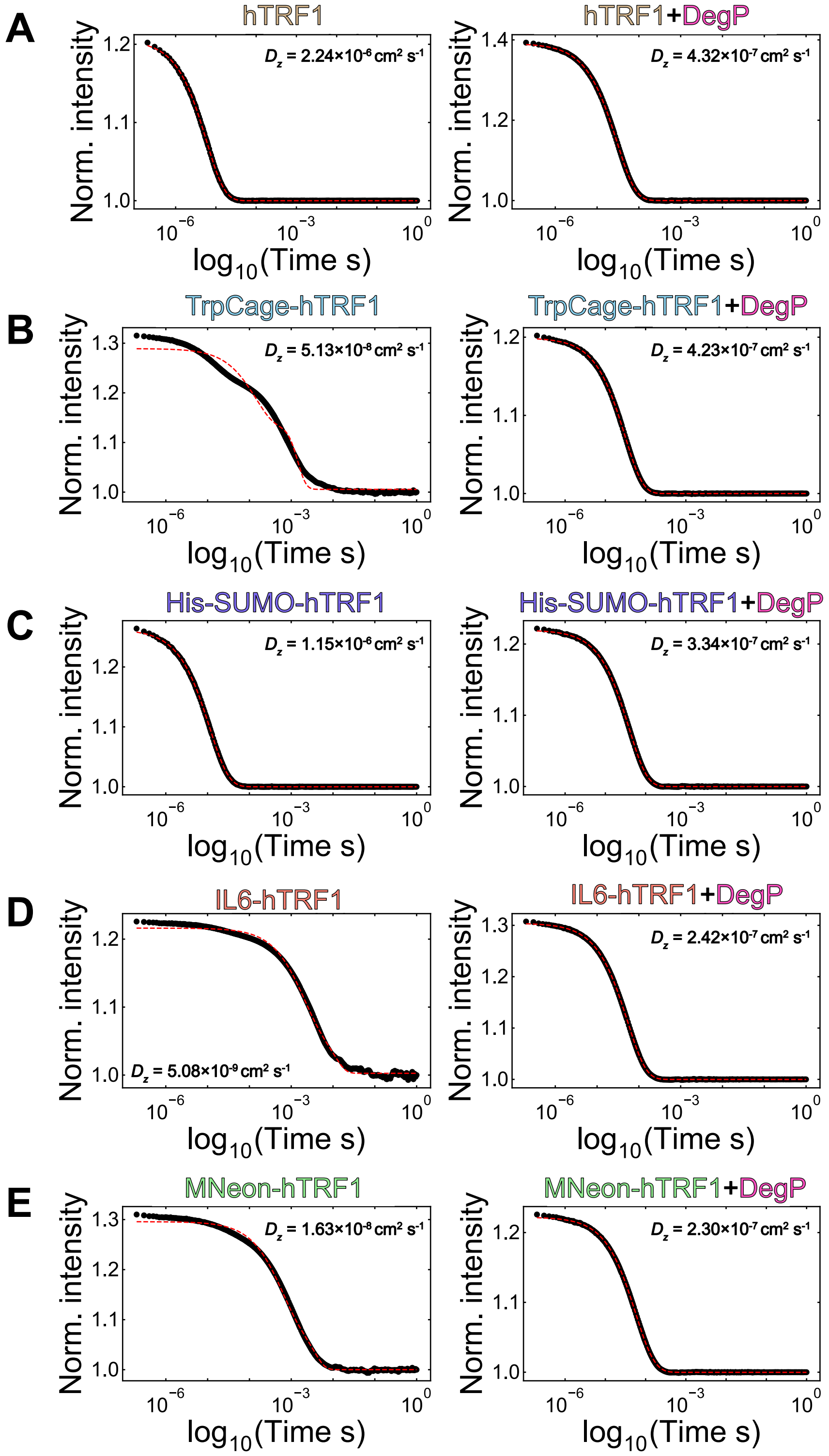

**Figure S1.** DLS autocorrelation data (black) for chimeric clients in the absence and presence of S210A DegP at 40 °C. (A-E, left) Autocorrelation data for apo hTRF1 (1200 μM), TrpCage-hTRF1 (1300 μM), His-SUMO-hTRF1 (250 μM), IL6-hTRF1 (300 μM), and MNeon-hTRF1 (400 μM) respectively. (A-E, right) Autocorrelation data for the substrates in (A-E, left) at 200 μM concentration bound to S210A DegP (100 μM). Data were numerically fitted (red) using in-house Python scripts to extract $D_z$ values (1). The poor fits for the TrpCage-hTRF1, IL6-hTRF1, and MNeon-hTRF1 datasets in (A-E, left) are the result of sample aggregation that occurs at elevated temperatures, as discussed in the main text. Higher protein concentrations were employed for the free clients in (A-E, left) in order to measure accurate $D_z$ values.

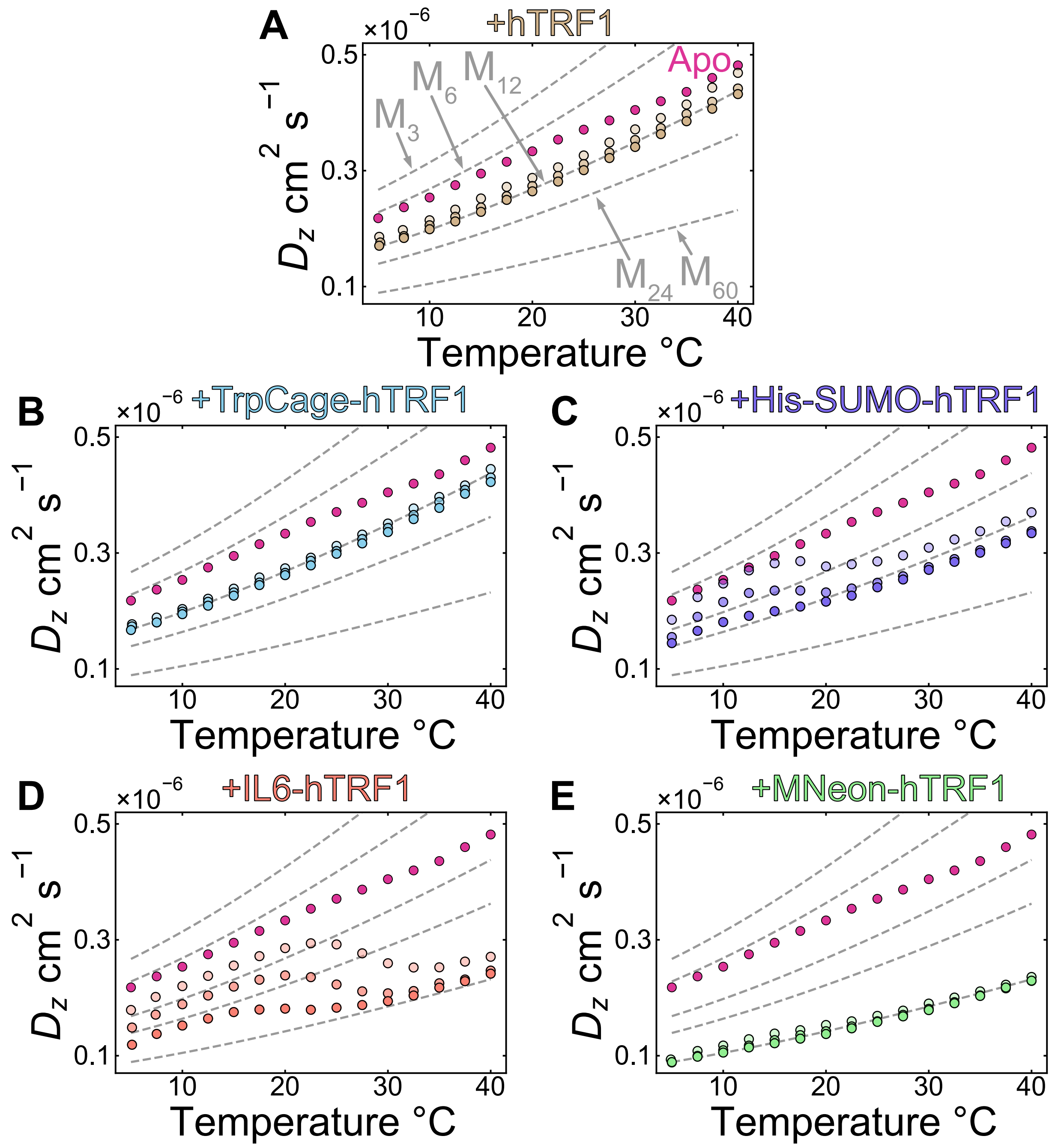


**Figure S2.** Characterizing DegP cages formed in the presence of the chimeric clients by DLS. (A-E) $D_z$ values for 100 μM S210A DegP measured in the presence of 50, 100, and 200 μM client proteins (light to dark colored circles respectively). The client protein investigated in each experiment is indicated above the plots. $D_z$ values for apo S210A DegP (red circles) are shown as a reference. $D_0$ values (grey dashed lines) calculated for 3mer, 6mer, 12mer, 24mer, and 60mer

DegP particles are indicated. In the case of the 3mer, 6mer, and 12mer, $r_h$s of 4.9, 5.7, and 7.8 nm, respectively, were obtained from $D_z$ values measured at low temperature (1), and then used to generate the $D_0$(T) profiles shown via the Stokes-Einstein equation. Profiles for the 24mer and 60mer oligomers where established from values of $r_h$s, 9.4 and 14.7 nm, respectively, as estimated at 20 °C using the structures determined here and the program HYDROPRO (12).

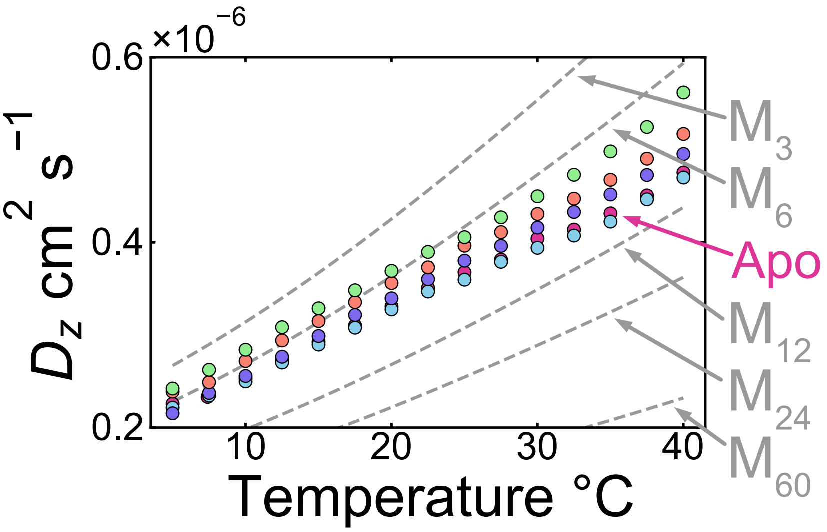


**Figure S3.** DLS analysis of DegP in the presence of the N-terminal portions of the chimeric clients. $D_z$ values for apo S210A DegP, S210A DegP + TrpCage, S210A DegP + His-SUMO, S210A DegP + IL6, and S210A DegP + MNeon are shown as red, blue, purple, orange, and green circles respectively. Grey dashed lines corresponding to the diffusion constants for 3mer, 6mer, 12mer, 24mer, and 60mer particles have been simulated according to Figure 2 and *SI Appendix*, Figure S2. These data were recorded with a 100 µM S210A DegP protomer concentration and 200 µM of each N-terminal fragment of the chimeric clients. Since $D_z$ values are intensity-weighted averages over each scatterer in solution (weighted by the molecular weight squared of each particle), in the absence of large cage formation, the smaller, unbound proteins in each well with DegP are expected to induce an upward shift in the $D_z$ values. The $D_z$ values recorded in the presence of these proteins reproduce the shape of the apo DegP profile (red circles) and are vertically shifted upward, indicating that no S210A DegP cages are formed. Note that the extent of the upward shifts increase as the size of the unbound clients grow, reflecting the molecular weight squared averaging of $D_z$.

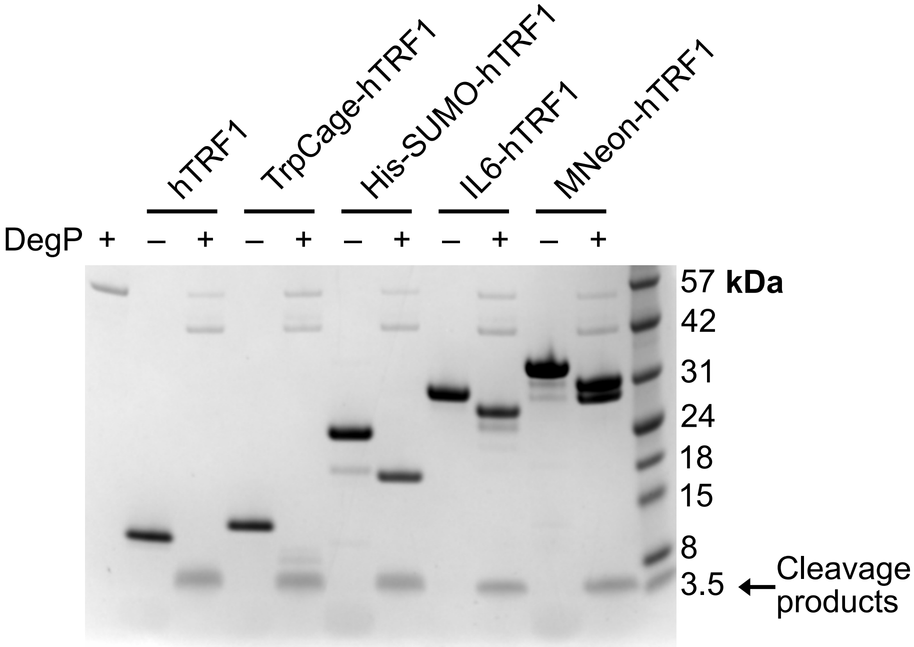


**Figure S4.** Cleavage of hTRF1 and chimeric clients by DegP assayed by SDS-PAGE. DegP undergoes a slow autocleavage reaction, as evidenced by the band at ~42 kDa in the substrate lanes +DegP. Client cleavage products from the reactions in the presence of DegP are indicated by the arrow next to the molecular weight marker at ~3.5 kDa.

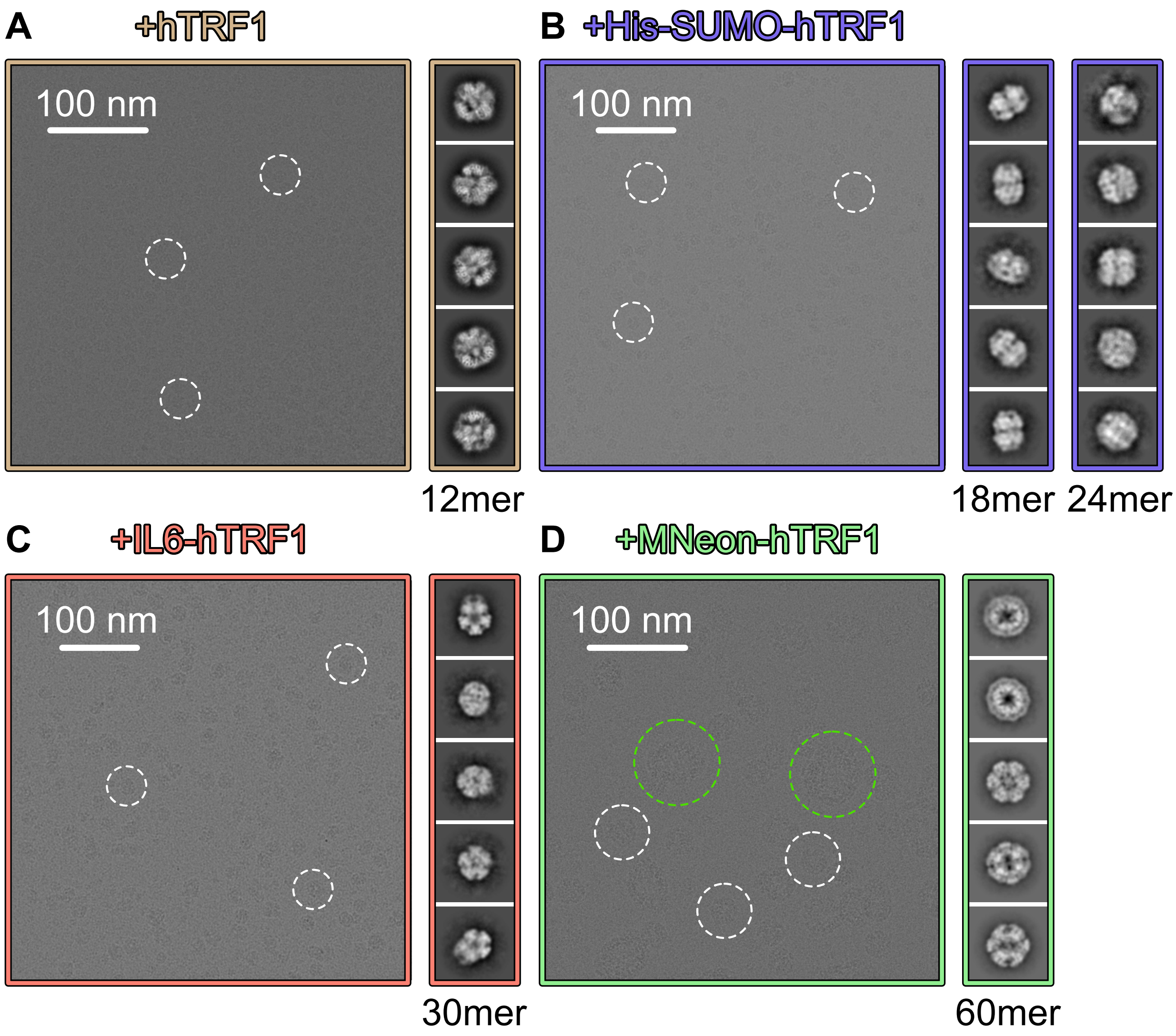


**Figure S5.** Micrographs and 2D class averages of DegP complexes that form in response to the engagement of the client proteins. Representative micrographs are shown for DegP cages that form in the presence of hTRF1 (A), His-SUMO-hTRF1 (B), IL6-hTRF1 (C), and MNeon-hTRF1 (D). 2D class averages for the DegP cage models that were constructed herein are shown to the right of the representative micrograph of the sample from which they were derived. Select particles within the micrographs are enclosed in white dashed circles. In (D), some of the extremely large DegP cages that form in the presence of MNeon-hTRF1 are encircled in green. A micrograph for DegP cages adopted in the presence of the TrpCage-hTRF1 client is not shown as these were found to be nearly identical to those for the hTRF1 client in (A), except with a small fraction of 18mer particles (see main text).

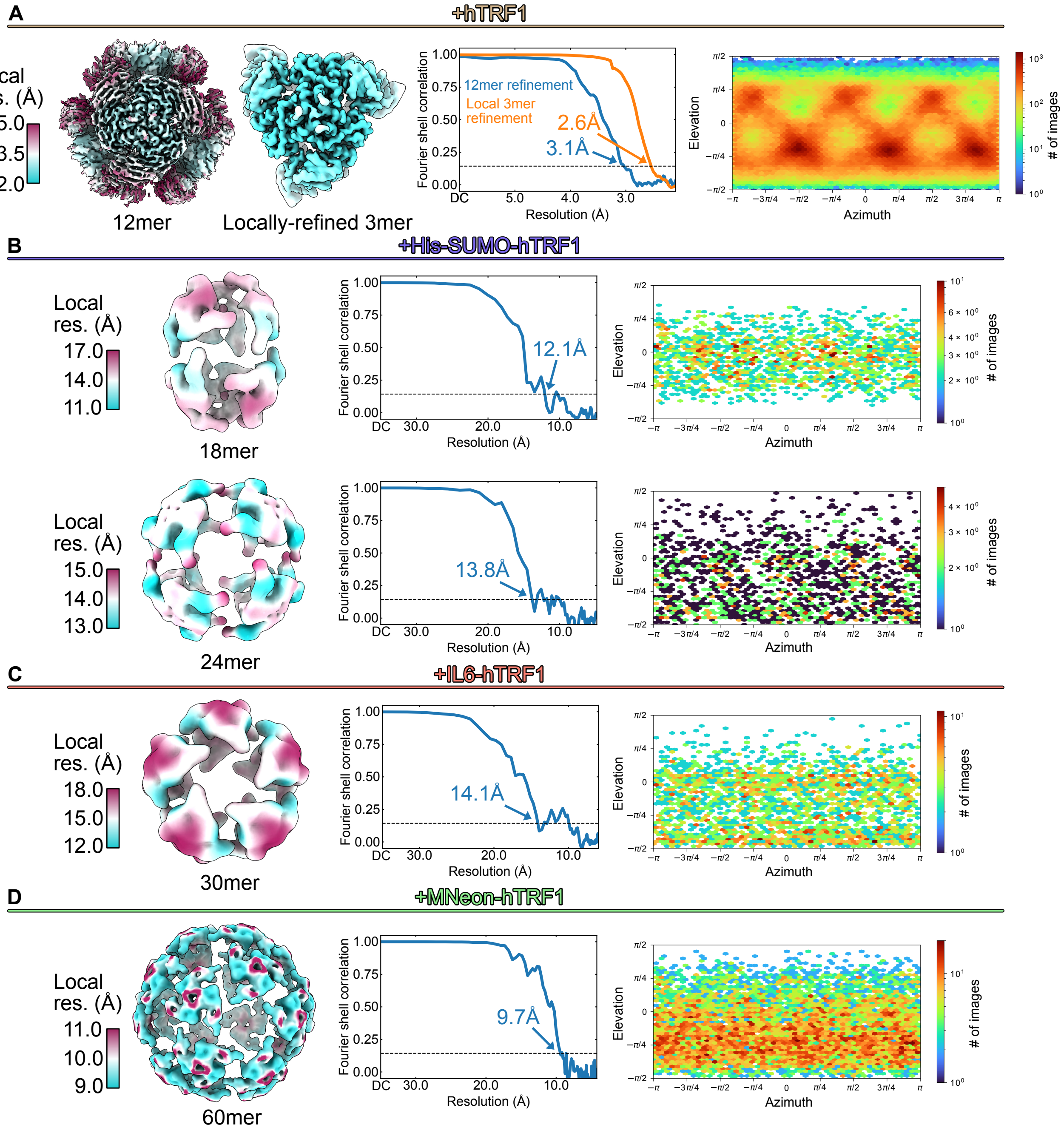


**Figure S6.** Cryo-EM maps of DegP cages adopted in the presence of clients. (A-D, left) Maps colored according to the local resolution for the 12mer and locally-refined trimer (hTRF1), 18mer and 24mer (His-SUMO-hTRF1), 30mer (IL6-hTRF1), and 60mer (MNeon-hTRF1) cages. (A-D, middle) Fourier shell correlation plots with global map resolution values. (A-D, right) Viewing direction distribution plots from the respective refinements in (A-D, left).

**Table S1.** Client protein amino acid sequences. The N-terminal domains of the chimeric clients are shown in black and their C-terminal hTRF1 tags are indicated in red.

| Construct | Amino acid sequence |
|---|---|
| hTRF1 | GRKRQAWLWEEDKNLRSGVRKYGEGNWSKILLHYKFNNRT<br>SVMLKDRWRTMKKL |
| TrpCage-hTRF1 | GNLYIQWLKDGGPSSGRPPPSGRKRQAWLWEEDKNLRSGVRKYGEGNWSK<br>ILLHYKFNNRTSVMLKDRWRTMKKL |
| His-SUMO-hTRF1 | MGSSHHHHHHSSGSDSEVNQEAKPEVKPEVKPETHINLKVSDGSSEIFFK<br>IKKTTPLRRLMEAFAKRQGKEMDSLRFLYDGIRIQADQTPEDLDMEDNDI<br>IEAHREQIGGGRKRQAWLWEEDKNLRSGVRKYGEGNWSKILLHYKFNNRT<br>SVMLKDRWRTMKKL |
| IL6-hTRF1 | GAPVPPGEDSKDVAAPHRQPLTSSERIDKQIRYILDGISALRKETCNKSN<br>MCESSKEALAENNLNLPKMAEKDGCFQSGFNEETCLVKIITGLLEFEVYL<br>EYLQNRFESSEEQARAVQMSTKVLIQFLQKKAKNLDAITTPDPTTNASLL<br>TKLQAQNQWLQDMTTHLILRSFKEFLQSSLRALRQMRKRQAWLWEEDKNL<br>RSGVRKYGEGNWSKILLHYKFNNRTSVMLKDRWRTMKKL |
| MNeon-hTRF1 | GVSKGEEDNMASLPATHELHIFGSINGVDFDMVGQGTGNPNDGYEELNLK<br>STKGDLQFSPWILVPHIGYGFHQYLPYPDGMSPFQAAMVDGSGYQVHRTM<br>QFEDGASLTVNYRYTYEGSHIKGEAQVKGTGFPADGPVMTNSLTAADWCR<br>SKKTYPNDKTIISTFKWSYTTGNGKRYRSTARTTYTFAKPMAANYLKNQP<br>MYVFRKTELKHSKTELNFKEWQKAFTDVMGMDELYKRKRQAWLWEEDKNL<br>RSGVRKYGEGNWSKILLHYKFNNRTSVMLKDRWRTMKKL |

**Table S2.** Peptide masses and their corresponding sequences obtained from LC-MS analyses of cleavage products of hTRF1 and chimeric clients after proteolysis by active DegP. Masses were identified using the FindPept tool (https://web.expasy.org/findpept/) assuming a mass tolerance of 1.5 Da. The intact mass of hTRF1 and its sequence are shown as a reference. Cleaved products are shown with their flanking residues in the intact hTRF1 chain in brackets. Note that for certain products, more than one fragment of hTRF1 can give rise to the observed mass at this level of detail (MS-MS is required for unambiguous sequence assignment) and thus multiple possible fragments are indicated in these cases. Since DegP processively digests the hTRF1 tag into many small fragments, as observed for other clients (13), only eight representative cleavage products are shown.

| Mass (Dalton) | hTRF1 sequence(s) |
|---|---|
| 6708.6 | GRKRQAWLWEEDKNLRSGVRKYGEGNWSKILLHYKFNNRT SVMLKDRWRTMKKL |
| 1078.5 | (L)HYKFNNRT(S) |
| 1264.6 | (L)HYKFNNRTSV(M) |
| 1304.7 | (I)LLHYKFNNRT(S) |
| 1490.8 | (I)LLHYKFNNRTSV(M) |
| 1492.8 | (V)MLKDRWRTMKK(L)<br>(S)GVRKYGEGNWSKI(L) |
| 1604.9 | (V)MLKDRWRTMKKL<br>(S)GVRKYGEGNWSKIL(L) |
| 1791.0 | (T)SVMLKDRWRTMKKL |
| 2949.5 | (G)EGNWSKILLHYKFNNRTSVMLKDR(W)<br>(G)NWSKILLHYKFNNRTSVMLKDRW(R)<br>(A)WLWEEDKNLRSGVRKYGEGNWSKI(L)<br>(S)KILLHYKFNNRTSVMLKDRWRTM(K)<br>(K)ILLHYKFNNRTSVMLKDRWRTMK(K) |

**Table S3.** Cryo-EM data acquisition and image processing.

| **TrpCage-hTRF1, His-SUMO-hTRF1 and IL6-hTRF1 Datasets** | |
|---|---|
| **Data Collection** | |
| Electron Microscope | Tecnai F20 |
| Camera | Gatan K2 summit direct detector |
| Voltage (kV) | 200 |
| Nominal Magnification | 25,000 |
| Calibrated physical pixel size (Å) | 1.45 |
| Total exposure ($e^-$/Å$^2$) | ~35 |
| Exposure rate ($e^-$/pixel/s) | ~5 |
| Number of frames | 30 |

| | | | |
|---|---|---|---|
| Defocus range (μm) | 1 to 3 | | |
| **Image Processing** | | | |
| Motion correction software | *MotionCor2* | | |
| CTF estimation software | *cryoSPARC v3* | | |
| Particle selection software | | | |
| 3D map classification and refinement software | | | |
| Sample | **TrpCage-hTRF1** | **His-SUMO-hTRF1** | **IL6-hTRF1** |
| Micrographs used | 166* | 119 | 189 |
| Particle images selected | 41,132* | 35,366 | 7,417 |
| **hTRF1 and hTRF1-MNeon Datasets** | | | |
| **Data Collection** | | | |
| Electron Microscope | Titan Krios | | |
| Camera | Falcon 4 | | |
| Voltage (kV) | 300 | | |
| Nominal Magnification | 75,000 | | |
| Calibrated physical pixel size (Å) | 1.03 | | |
| Total exposure ($e^-/Å^2$) | ~45 | | |
| Exposure rate ($e^-$/pixel/s) | ~5.5 | | |
| Number of frames | 29 | | |
| Defocus range (μm) | 0.9 to 2 | | |
| **Image Processing** | | | |
| Motion correction software | *MotionCor2* | | |
| CTF estimation software | *cryoSPARC v3* | | |
| Particle selection software | | | |
| 3D map classification and refinement software | | | |
| Sample | **hTRF1** | **hTRF1-MNeon** | |
| Micrographs used | 4,383 | 1,185 | |
| Particle images selected | 844,585 | 14,443 | |

*For the DegP:TrpCage-hTRF1 sample, the motion corrected micrographs were not subjected to further analysis beyond particle population estimates as they were determined to be mainly 12mer.

**Table S4.** Cryo-EM map and atomic model statistics.

| **Dataset** | **DegP:hTRF1 3mer local refinement** |
|---|---|
| **Modelling and refinement software** | *Coot*, *Rosetta* |
| **Protein residues** | 1212 |
| **Ligand** | -- |

| | |
|---|---|
| **RMSD bond length (Å)** | 0.031 |
| **RMSD bond angle (°)** | 1.738 |
| **Ramachandran outliers (%)** | 0 |
| **Ramachandran favoured (%)** | 99.24 |
| **Rotamer outliers (%)** | 0 |
| **Clash score** | 0.44 |
| **MolProbability score** | 0.66 |
| **EMringer score** | 4.6 |
